# How does an ectodomain of membrane-associated proteins stand upright and exert robust signal?

**DOI:** 10.1101/2020.07.29.226837

**Authors:** Swetha Lankipalli, Udupi A. Ramagopal

## Abstract

Even after decades of research, a comprehensive mechanism that elucidates the underpinnings of signaling through the cell membrane is still elusive. Here, we address a simple question- “how does the ectodomain of a membrane-associated protein consisting of multiple domains and connected by flexible linkers stand ‘upright’ on the membrane?”. Our analysis based on large amount of available structural and functional data, looking for a pattern of association of these molecules in the crystal structures and with the concept that ‘random things seldom repeat’ lead to a surprisingly interesting and consistent observation that (1) the weak *cis-*interaction mediated symmetric oligomerization of signaling molecules not only support their ‘upright’ orientation but often bury their ligand-binding surface to avoid spurious signaling (2) the linkers connecting the domains are probably not flexible as presumed. This analysis provides a model for pre-liganded receptor supramolecular organization that resolves some of the mysteries unanswered by hypothesis such as ‘lipid-rafts’ and ‘fence and pickets. With CD4, pMHCII, CD2 and TNFR1 as examples, we show that the observed *cis-*association of molecules also correlate well with their functional role. Further, our analysis reconciles the long-standing controversies related to these molecules and appear to be generic enough to be applied to other signaling molecules.

## Introduction

Membrane-associated proteins constitute ~50% of the total cell-surface volume and play a critical role in the detection and integration of extracellular mechanical, physical and chemical stimuli and hence play a vital role in cell-cell recognition and communication[1]. Its organization is of interest for over a century. Nano, meso, and micro-clusters of membrane-associated proteins have been revealed by super-resolution microscopy**(see, supplementary Information-section 2.1)** and now can be quantitatively investigated[2]. It is also known that on the cell membrane, these proteins cluster and organize/reorganize depending on the functional context such as polarity, adhesion, immune, and synaptic functions [3–5] However, what is not clear is; how do they achieve this compartmentalization and how the clustering state imparts the ability to perform the accurate and reproducible function? Models such as ‘lipid rafts’ [6] molecular crowding [7] and ‘fences and pickets’ have been proposed [8] to explain the clustering phenomenon. Unfortunately, the diversity of these clusters can’t be explained satisfactorily by any of these models [5, 9–15]. Further, the control of association of these proteins with large sequence and architectural diversity, by either lipid-rafts or by the intracellular cytoskeleton network is difficult to perceive. For example, most clusters are homotypic [16, 17] which requires a mechanism that can segregate proteins based on their identity. Recent reports propose that the ability of these proteins to cluster with each other (homo or hetero depending on the cluster) is inherent in them and is mediated by weak protein-protein interactions[9, 18, 19]. However, the physicochemical signatures governing the sorting of these molecules in to clusters of different sizes and architecture, with the ability to reorganize and perform reproducible and accurate function avoiding spurious signaling remain unanswered[4].

Interestingly, even a much fundamental question, how cell-surface proteins position themselves ‘upright’ on the membrane prior to their interaction with a cognate partner on the opposing cell is a mystery. The ectodomains of these proteins vary in size, architecture, and most of them consist of multiple domains connected by linkers (Fig. 1). Further, they are membrane-tethered via an extended extracellular juxtamembrane linker (hereafter called as EJ-linker) which is purported to be flexible. For these membrane proteins to exert reproducible organization and function, they are expected to be robust and at the same time pliable to respond to weak cues and transform them into robust signals across the membrane. It is intriguing to know, how these considerably large cell surface molecules connected by multiple interdomain linkers maintain appropriate orientation and perform an accurate function. Further, flexible linkers can increase the conformational entropy of a protein on the cell membrane, and hence it is likely that such molecules on the crowded membrane are prone to tangle with each other forming aggregates that can never respond to a specific signal.

**Fig.1.**
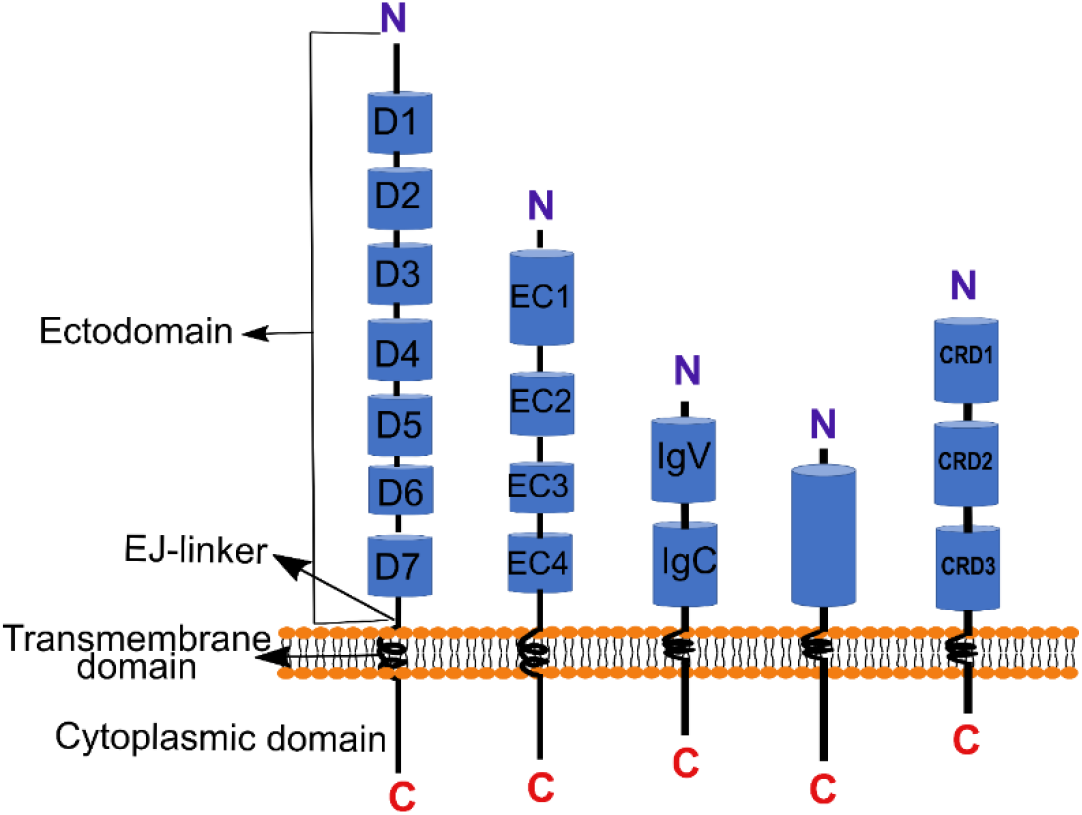
Representative structures of diverse type-I membrane associated proteins. The naming convention used are also diverse-they are named as D_1_ to D_n_ (Domain 1 to Domain ‘n’), EC1 to EC_n_ (ectodomain 1 to ‘n’), CRD1 (Cysteine rich domain 1 to ‘n’), where is ‘n’ is an integer and also sometimes they are named based on their fold such as IgV, IgC_1_, IgC_2_, etc. The multidomain proteins with a stalk like EJ-linker (extracellular juxtamembrane linker) are tethered to the membrane by a single pass transmembrane helix. The N terminus is at the membrane distal end on the extracellular side while the C terminus is on the cytoplasmic side of each protein.

In most of these cases, the extracellular domains distal to the membrane are the ones that interact with their cognate partners on the opposing cell. Hence, these molecules are expected to stand ‘upright’ on the cell surface and position the membrane-distal domain(s) in an optimal orientation on the cell surface to ensure efficient function. Further, the elegant ‘kinetic segregation’ model that explains the segregation of molecules in the cell-cell contact during T-cell signaling requires that the distance spanned by each molecule from the membrane is proportional to the effective length of the molecule [20]. Although, current techniques such as super-resolution microscopy [21], single-molecule tracking [22, 23]and other allied techniques have revolutionized the way we look at the biological systems[24], unfortunately, even the improved resolution does not provide a comprehensive mechanism for controlling the association and function of these molecules[25].

We asked the question, whether the packing analysis (association in the crystals) of pool of available crystal structures provide any clue on preferential and organized association and whether such an association (though weak) manifests itself in the crystal, revealing physiologically relevant pre-liganded *cis* interactions? In fact, the insights that can be gained from crystal structure analysis in the case of adhesion molecules has been extensively discussed by Honig. B and Shapiro. L[26] **(For more such examples see, supplementary Information-section 2.2)** Since the crystal formation requires epitope(s) of association[27] on the surface of each molecule that can support the formation of 3D lattice, in the absence of any preferred interaction, these molecules are expected to have an infinite modes of self-association. In contrast, if the ectodomains of these proteins on the same cell have physiologically relevant pre-liganded association preference (*cis* association), although weak-to-be-detected in solution, we pondered whether any such preferred association can be observed in the crystals from the pool of available crystal structures of these receptors. The proteins described here are those that fit into the following criteria (1) ectodomains that are known to be monomers in solution (or oligomers weak-to-be-detectable in solution) (2) multiple crystal structures have been determined for the complete ectodomain of the same molecule either from the same organism or from different organisms. This criterion was set to make sure that the observed association in the crystal is not an accidental observation and those that repeat might have some significance (random things seldom repeat) (3) since our question is regarding the pre-liganded association, to eliminate the effect of cognate binding partners, only apo structures were considered first and later they were compared with the complex structures containing cognate partner (4) we did not consider homomers such as cadherins and other cell-adhesion molecules as they tend to organize with stronger (detectable) *trans* and *cis* interactions in the crystals. The *cis* association observed in the crystals of adhesion molecules are result of post *trans* association and hence may not represent *cis* association of these molecules on the individual cell.

Based on our analysis, we propose that the cryptic *cis-*oligomerization and supramolecular association of rigid ectodomains together with conformational entropy constrained by membrane tethering helps them to stand ‘upright’ on the cell surface. More interestingly, these weak oligomers often hide their ligand-binding surface to avoid spurious signaling and can further associate into higher-ordered arrays. Importantly, this analysis reconciles some of the contradicting observations pertinent to T-cell signaling mediated by MHCII-TCR-CD4 and other molecules such as CD2 and TNFRs involved in immune signaling.

## Results and discussion

### Interdomain flexibility and the role of EJ-linker

To understand the interdomain flexibility of multi-domain cell-surface receptors, the crystal structures of several of these molecules present on immune cells were analysed. The structure of receptors containing 2-3 Ig domains, such as CD2, NTB-A, PD-L1, PD-L2, B7-1, Nectin1, Nectin2, Nectin3 and so on reveal that the interdomain hydrophobic interactions help in maintaining a stable relative orientation [28–33] Similarly, in the case of larger molecules such as CD4, cadherins, ephrins, CD45, CD22 and so on, it has been observed that the structure of truncated proteins with few domains superposes well with the complete ectodomain structures of these proteins**(Fig. S1)** [34–38] In molecules such as cadherins, CEACAM-1, Syntaxin, SNAP25 and so on, the interdomain stabilization is mediated by calcium ions[39–41] **(supplementary information- section 3.1)**. Also, in most of these proteins, solution studies confirm that the structures observed in the crystals are highly similar to their conformation in solution [36, 40]. The above analysis suggests that these receptors containing multiple domains connected by linkers are rigid or posses’ minimal inter-domain flexibility. The observed interdomain stability of the ectodomain cannot be a complete answer for their ability to stand ‘upright’ as they are tethered to a jelly-like membrane via an EJ-linker which is presumed to be flexible. Since the EJ-linker is in the aqueous environment just outside the cell surface, if flexible, it is expected to contain more of hydrophilic residues. Our analysis suggests that it is not always true. For example, in some of the key molecules involved in immune function, the sequence of the EJ-linker contains close to 50% hydrophobic residues and sometimes rich in conformation constraining residues like proline, together with aromatic residues **(Table-1).** Such sequences are expected to be rigid and being substantially hydrophobic, their existence in the aqueous extracellular region requires burial of these hydrophobic residues.

**Table-1.**
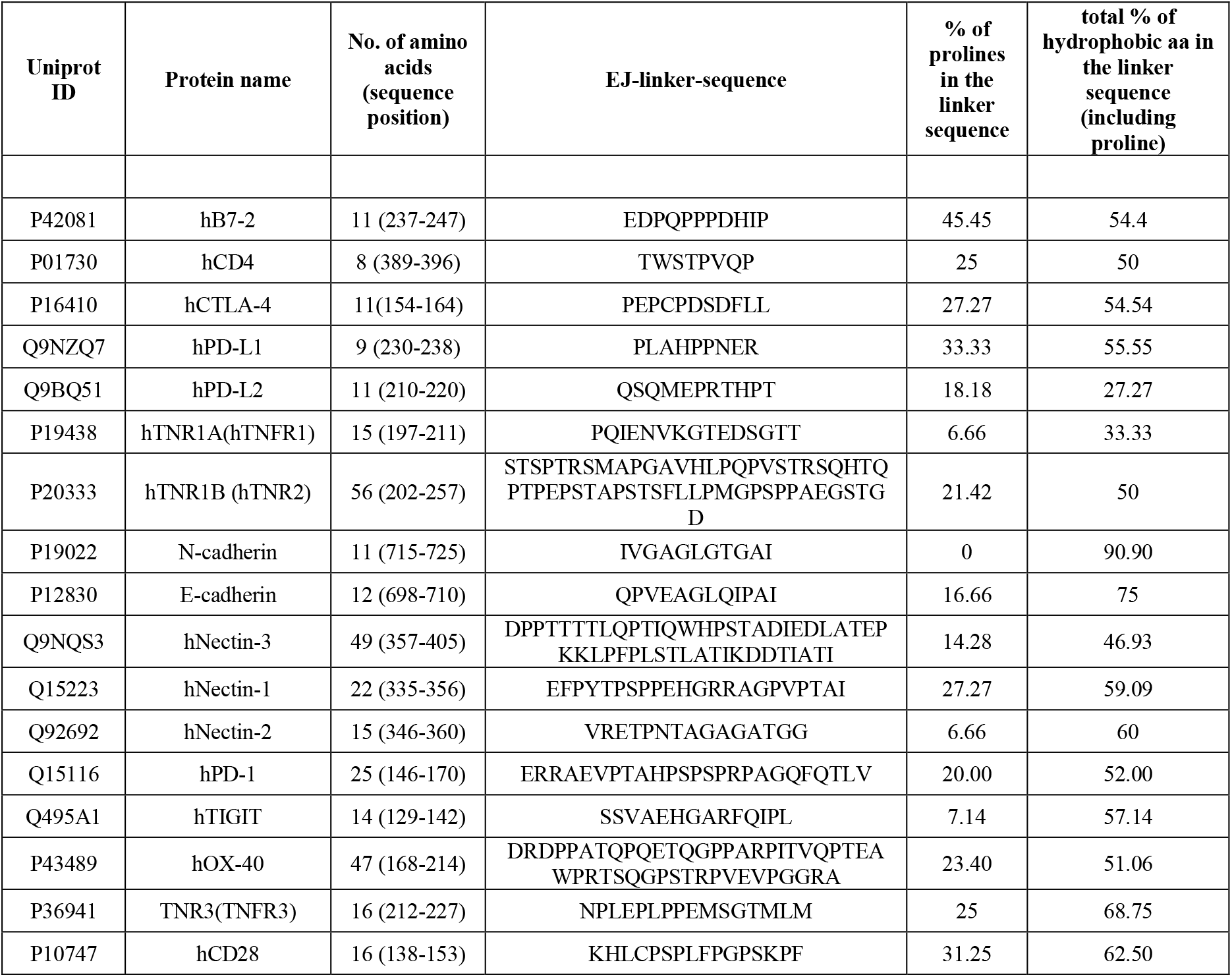
Proline rich sequences of extracellular juxtamembrane linker(EJ-linker) of selected type-I membrane associated proteins and overall percentage of hydrophobic residues.

This raises the question of whether they are flexible as generally presumed or such linkers from adjacent molecules on the cell membrane can associate with each other through their hydrophobic residues, stabilizing well-defined oligomers and/or have the ability to get involved in energetically similar but different oligomerization states? Unfortunately, EJ-linker is the most neglected part of all the studies directed towards structural characterization, oligomerization, and clustering. In most such studies involving cell-surface molecules, manipulations are done on the ectodomain (excluding the EJ-linker) and/or on the intracellular domain. **(see supplementary information section 3.2 for few specific examples)** These observations suggest that the inclusion of the EJ-linker region is important in studies involving, oligomerization and signaling. However, there are few examples, where EJ-linker oligomerization has been shown to impact the signalling ability of receptors. In the following section, we briefly discuss few examples where importance of juxtamembrane linker has been observed.

Tropomyosin receptor kinases (Trks- TrkA, TrkB and TrkC) which also belongs to the family of receptor tyrosine kinases (RTKs) are important receptors in the development of nervous system and are activated by different nerve growth factors (NGFs). The architecture of a typical Trk has an extracellular domain with a membrane distal Leucine rich domain (D1), followed by two cysteine rich domains (D2-D3) and two Ig like domains (D4 and D5). The D5 domain is followed by an extra-cellular juxtamembrane linker (EJ-linker), a transmembrane domain and a cytoplasmic kinase domain. Dimeric ligand (NGF) binds to the D5 domain of Trk and like other RTKs, Trks were initially considered to form ligand induced dimers. In contrast, many biochemical studies such as chemical cross-linking experiments, protein fragment complementation assays and crystal studies of the kinase domain have shown that Trks could exist as pre-formed inactive dimers[42]. Also, the study has shown that the pre-formed inactive dimers are mediated by interactions of EJ-linker and by deletion of EJ-linker, it was shown that EJ-linker acted as an inhibitory motif for the formation of active dimer. The binding of ligand to the D5 domain was presumed to over-ride the inhibitory interactions of EJ-liker leading to ligand induced active dimer formation. Further, Franco et.al., have supported the role of EJ-linker in the pre-liganded inactive dimer by mutation studies, MD simulations and have also elucidated the dimerization of transmembrane domain by NMR studies concluding that EJ-linker and transmembrane domain are involved in the pre-liganded inactive dimerization of Trks[43]. Similarly, EGFR is one of the well-studied receptor tyrosine kinase. Replacement of juxtamembrane-linker with an unstructured (GGS)10 sequence abolished phosphorylation of all the tyrosine residues, affected receptor dimerization and ligand binding capability. There are several similar studies showing the effect of EJ-linker on receptor functions, particularly on the receptor tyrosine kinases[44]. In the case of T-cell co-inhibitory receptor CTLA-4 (cytotoxic T lymphocyte-associated protein-4). It is a CD28 family member and the ligand binding domain of CTLA-4 is a weak homodimer[45]. Various studies have shown that CTLA-4 forms a stable dimer only in the presence of EJ-linker (with sequence ^119^Pro-Glu-Pro-Cys-Pro-Asp^124^ which is also conserved in all CD28 family members). Consistent with this, in the structure of CTLA-4 that includes proline rich EJ-linker, it has been observed that the linker together with few hydrophobic residues results in a stable dimer which is further stabilized by a disulfide bond between the two protomers of the dimer[46].

Apart from the putative role of EJ-linker and limited inter-domain flexibility of ectodomain, the preferential association of ectodomains of adjacent identical proteins (*cis-*interactions) appears to be important for ‘upright’ (or appropriate) orientation and robust signaling. In the following section, we present few cases, where ectodomain of these receptors are known to be monomers in solution (or more accurately, possess undetectable oligomerization potential), but consistently exist as multimers in the crystal, maintaining similar physiologically relevant association. Other than the criteria used for the selection of molecules discussed above, we restricted ourselves to immune molecules, the field of our expertise, where we can effectively correlate their function to the observed association preference.

### 1D array of CD4

The CD4 is a key co-receptor on the T-cells that interact with peptide-loaded major histocompatibility complex-II (pMHCII) on the antigen-presenting cells (APCs) and contributes to signal transduction through its cytoplasmic tail by interacting with lymphocyte kinase (Lck) [47]. The ectodomain of hCD4 consists of four Ig domains (named D1 to D4), where the membrane distal domain (D1) is shown to interact with pMHCII[48]. Although there is no clear indication of dimerization of CD4 ectodomain in solution[49], the dimeric association of CD4, involving the membrane-proximal D4 domain has been observed in three different isoforms of CD4(D1-D4) crystals (PDB-entry: 1WIQ, 1WIP, 1WIO**; Fig. 2A)**[34]. The description of the structure in the author’s own words is very interesting “*the D4-D4 associated dimers are butterfly-shaped in profile, with D1-D3 as ‘wings’, D4 pair as ‘torso’ and C termini as ‘legs’, as if perching on the hypothetical membrane surface perpendicular to the dyad axis*”, which clearly depicts the dimeric ‘upright’ orientation of CD4 on the cell membrane. While analytical ultracentrifugation and light scattering studies did not support CD4 oligomerization, crosslinking, mutational and immunoprecipitation studies confirmed the role of the D4 domain in CD4 dimerization[34, 50]

**Fig.2.**
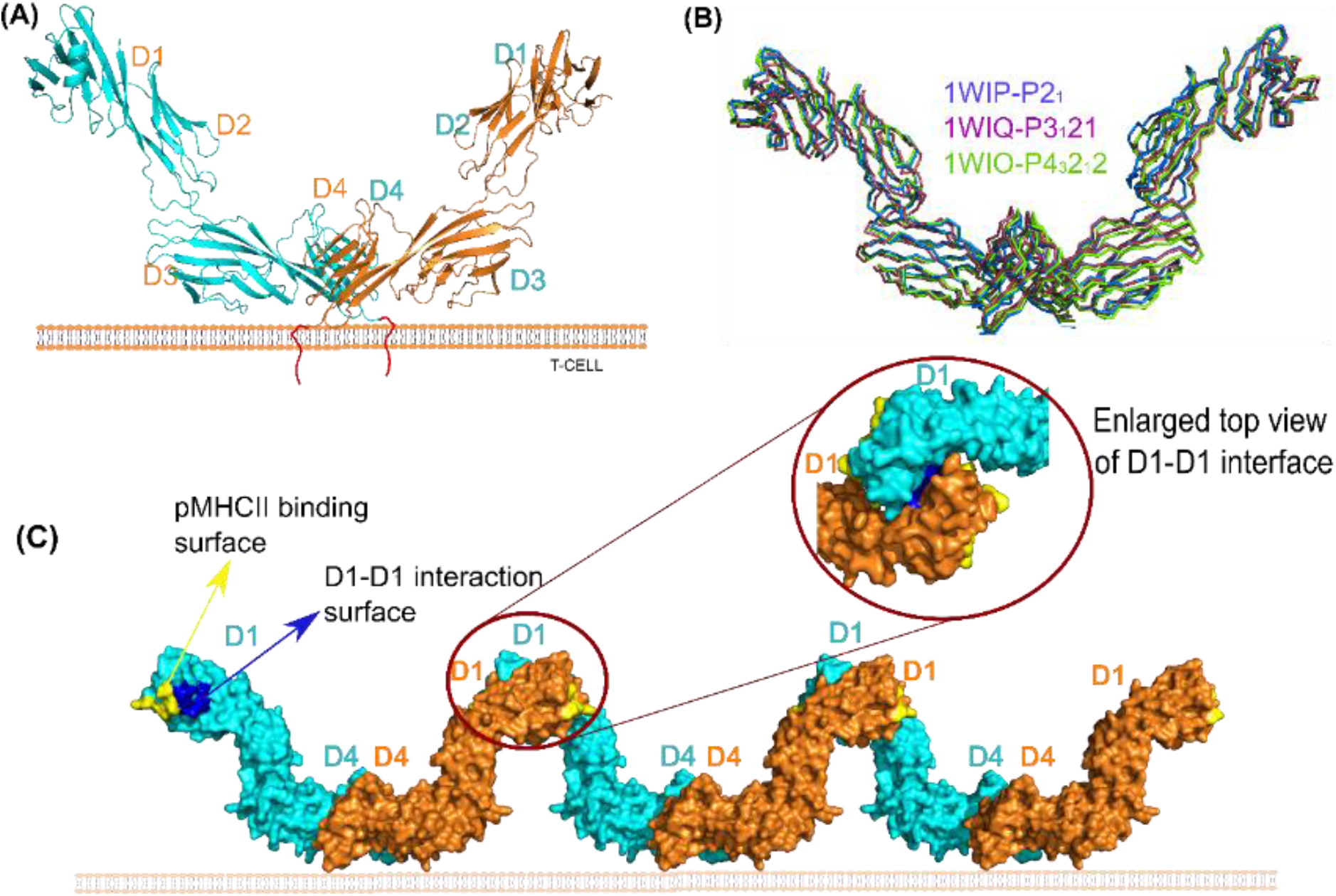
Structural details of ectodomain of CD4 (D1-D4) **(A)** Cartoon representation of dimeric CD4, the monomers coloured in cyan and orange associate via their membrane proximal D4 domains and the D4 domains interact with the membrane placing the D1-D3 domains distal to the cell surface. **(B)** Superposition of three different crystal structures of CD4 (ribbons) (PDB entries: 1WIP, 1WIQ, 1WIO) highlighting the exactly similar D4-D4 mediated dimeric association of two protomers (maximum Cα RMSD of ~1.2Å). **(C)** packing analysis of the three different structures of CD4 (surface representation) reveals a formation of identical 1D array of CD4 homodimers due to D4-D4 interaction and D1-D1 interaction conserved in all the three structures. The D1-D1 interface (shaded in dark blue), partially overlaps with pMHCII binding site (shaded in yellow) precluding its binding to MHCII in non-*signaling* resting state and hence appear to avoid spurious signaling.

Contrary to this observation, there is a considerable argument in favour of CD4 dimerization through the D1 domain[51] and independent experiments involving mutations on either D1 or D4 domains resulted in impaired dimerization[50, 52, 53]. In fact, our packing analysis of these molecules in the crystals of complete ectodomain of CD4 reveals that the pairing of D4-D4 domains together with the pairing of D1-D1 domains form a zigzagged 1D array of CD4 molecules **(Fig. 2C)** (similar to a portable zigzagged display panel). Consistent with this observation, at higher concentrations (5mg/ml) analytical ultracentrifugation studies revealed a non-ideal behavior of CD4 in solution[34]. Further, we also note that the D1-D1 domain interaction observed in the above structures is canonical which is generally observed in most receptor: ligand interaction involving IgV domains. Moreover, the same mode of interaction is conserved in all the six different crystal structures containing only D1-D2 domains, **(Fig. S2)**, and is in line with mutation studies mentioned above.

Therefore, (1) the conservation of similar association of D1 domains in all the nine crystal structures, (2) their canonical association (3) the functional data that corroborates well with observed D1 and D4 dimers, suggest the probable physiological role of dimerization of CD4 at both ends. Moreover, the structures of all the apo CD4 monomers, truncated structures, and the one that is observed in the MHCII:TCR:CD4 ternary complex[54] superpose well with each other, suggesting the elongated CD4 ectodomain is rigid. Hence, it appears that the rigid structure of CD4 ectodomain together with zigzagged 1D array, (a supramolecular organization of CD4) supports the erect orientation of CD4 molecules on the cell surface. The most interesting observation that emerged out of this analysis is that the membrane distal D1-D1 interface that appears to support the ‘upright’ orientation of CD4 molecules on the cell surface is also used for pMHCII binding, which closes itself in supramolecular organization. Hence, the formation of pre-liganded 1D array and consequent closing of pMHCII binding site on D1, not only helps its ‘upright’ orientation on the cell surface but also appear to avoid spurious interaction with the pMHCII molecules present on the opposing APCs during the screening of peptide by TCR. Consistent with these observations, in a study aimed at determining the association between HIV viral entry and CD4 receptor dimerization, both D1 mutants and D4 mutants were used. Here they report the increased HIV viral entry efficiency in the case of both the mutants compared to wt CD4. The reduced dimerization observed in both D1 mutants, as well as D4 mutants, suggested that the two different CD4 dimeric forms (D1 dimer as well as D4 dimer) co-exist on the cell surface[52]. As explained in the main-text, closing of D1 domain (that interacts with MHC-II as well as gp120 of HIV) is essential for avoiding spurious signaling. The continous array of CD4 shown in our analysis is prone to be disturbed by either of the mutants and hence, disturbance of dimerization probably results in uncontrolled interaction D1 domain with gp120 making its entry efficient.

### New model for pMHC-TCR-CD4 mediated signaling

The pMHCII-TCR-CD4 complex is central to immunological synapse formation. Although signaling through pMHCII with TCR and CD4 is important, the mechanism by which such an accurate process is achieved is still enigmatic. Recognition of pMHCII present on the APCs by TCR on T-helper cells is known to trigger the interaction of co-receptor CD4 (present on T-helper cells) with pMHCII. pMHCII is a heterodimer consisting of two chains named α and β, with the peptide-binding groove crafted from residues involving both the chains. The first report describing the two crystal structures of pMHCII (HLA-DR1 with antigenic peptide) from the human B-cell membrane describes a dimer of heterodimers (later termed as superdimers) in both the crystal forms(PDB entries: 1DLH, 1AQD)[55]. The parallel association of two heterodimers brings all the four C-termini in one direction with both the peptide binding groves facing in the opposite direction that supports the presentation of bound peptides in an orientation appropriate for recognition by TCRs on juxtaposed T-helper cells **(Fig. 3A).** Here, facing of the C-terminal end of all the four molecules in one direction and consequent tethering to the membrane at four different points in proximity appears to stabilize the ‘upright’ orientation of MHCII. The authors also propose that the superdimer might bind to CD4 coreceptor with increased affinity and hence initiate TCR mediated signaling. Subsequent determination of several crystal structures of pMHCII variants (HLA-DR1; PDB-entries: 1KG0, 4I5B, 1AQD) including HLA-DR1 in complex with superantigen *Staphylococcus aureus* enterotoxin B (PDB entry: 1SEB) and HLA-DR3 (PDB entry: 1A6A) revealed a similar association of dimer of αβ heterodimers. This superdimer is expected to be very weak as the interface is formed by discontinuous stretches of polar and charged residues with very few hydrophobic interactions [55]. This might probably be the reason for the lack of any evidence of such superdimer in solution as measured by hydrodynamic radii and other techniques[56, 57]. However, chemical crosslinking studies and immunoprecipitation of pMHCII dimers (I-Ek) from B-cell[58, 59] also supported the “superdimer mediated T-cell activation” hypothesis. Later, the structure determination of complete ectodomain of CD4 and the observation that the CD4 also exists as a dimer (discussed in the previous section on CD4), triggered interest in investigations on the possibility of dimeric CD4 and pMHCII superdimer mediated T-cell signaling[53].

**Fig.3.**
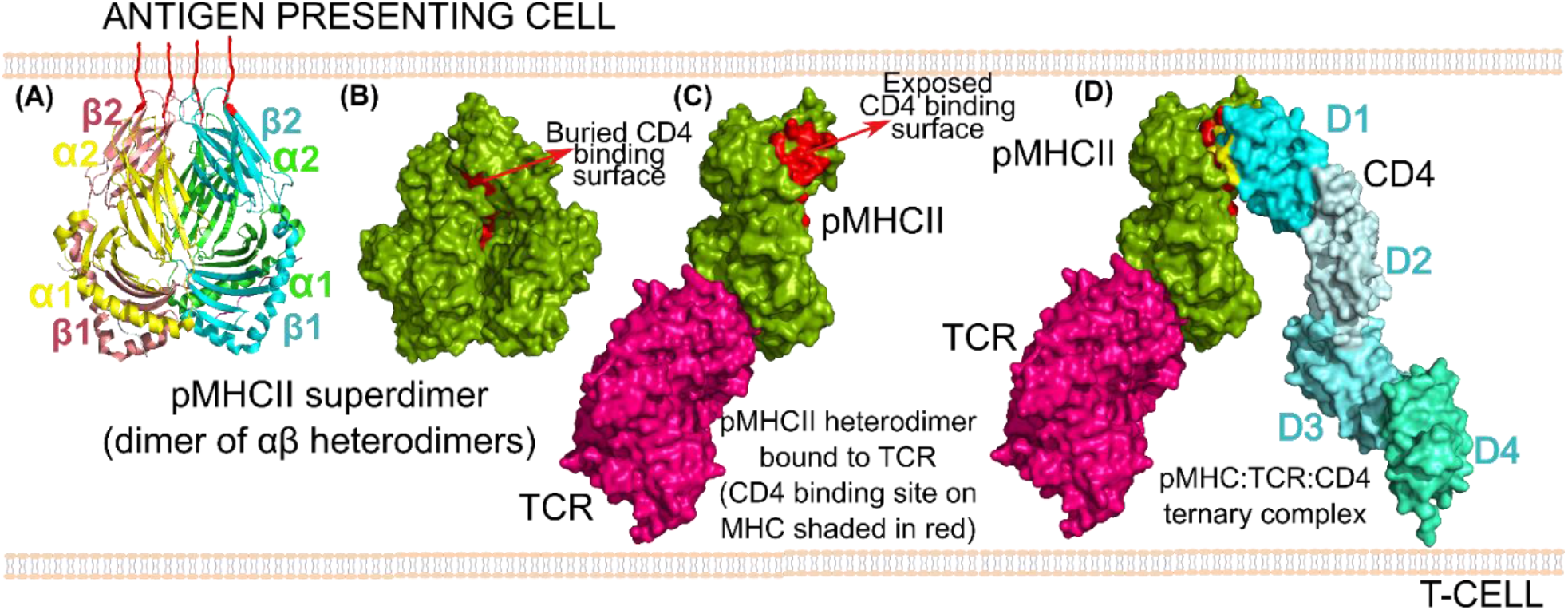
A new model of pMHCII:TCR:CD4 *signaling*: **(A)** Cartoon representation of pMHC-II superdimer observed in different crystal structures depicting dimer of heterodimer formed by an α and β chain. The C-termini of all the four chains (2α and 2β) face in the same direction as if emerging from the same cell. Thus, it is anchored at four points stabilising the *‘*upright*’* orientation **(B)** Surface representation of the pMHCII superdimer hiding the CD4 binding site (shaded in red) in the dimer interface **(C)** TCR (magenta):pMHCII (green) complex reveals that pMHCII exists as heterodimer but not as superdimer in the complex. Note that the disassembly of superdimer into heterodimer exposes the CD4 binding site (shaded in red) on MHC. **(D)** Ternary complex structure of pMHC:TCR:CD4(cyan) (PDB entry: 3T0E) showing simultaneous interaction of TCR and monomeric CD4 with pMHCII heterodimer. CD4 interacts with MHCII via its D1 domain.

Interestingly, the superdimer which was observed in most of the pMHCII variant crystal structures was not seen in any of the crystal structures of pMHCII in complex with TCR (PDB-entries: 1FYT, 2ICW, 2WBJ, 2XN9, 1ZGL, IJ8H, etc.,), in the structures of pMHCII in complex with CD4(D1-D2, PDB entries: 1JL4, 3S4S, 3S5L, etc.,) and also in the ternary complex structure of pMHCII-TCR and CD4 (PDB entry: 3T0E)[54]. The structures of pMHCII in complex with CD4 (D1-D2) revealed that the CD4 binding region is in the interface formed by α_2_ and β_2_ domains on pMHCII which is completely hidden in the pMHCII superdimer interface[48]. The authors also propose a V-shaped model for the TCR-MHC-CD4 ternary complex which later matched with the crystal structure of the ternary complex (PDB entry: 3T0E)[54], with complete ectodomain of CD4. With the absence of superdimeric structures of pMHCII in complex with TCR and/or CD4, the earlier models which emphasized pMHCII superdimer as a functional entity lost their relevance and it is now established that TCR-pMHCII-CD4 is a complex containing a single unit of each protein.

Unfortunately, the importance of such an elegant pMHCII superdimer which was observed in many crystal-structures of pMHC and proven to be present on the cell-surface by other studies [59]was comfortably forgotten thereafter. Based on the observations made from all the currently available crystal structures of pMHCII superdimers, pMHC-TCR structures (>25 structures), pMHC-CD4(D1-D2), and the ternary complex structures of TCR-pMHC-CD4, we provide a model that reconciles all the experimental observations and highlight the importance of pMHCII superdimer. As observed in the pMHCII structures, the superdimer might be the native state before binding to TCR which precludes CD4 binding, as the CD4 binding site is buried in the superdimer interface **(Fig. 3B).** When the MHCII loaded with antigenic peptide interacts with specific TCR and with an appropriate strength, the resulting mechanoforce might dislodge the weak pMHCII superdimer, exposing the CD4 binding site **(Fig. 3C).** As mentioned above, this is consistent with all the TCR: pMHCII crystal structures, where monomeric TCR is observed to interact with pMHCII heterodimer (and not the superdimer) with exposed CD4 binding site **(Fig. 3C)**. Subsequently, the binding of monomeric CD4 to pMHC-TCR complex forms the V-shaped arch **(Fig. 3D)** as observed in the ternary complex structures and probably results in downstream signaling initiated through Lck. Thus, it appears that pMHCII is an obligate superdimer that hides its CD4 biding site and avoids CD4 interaction prior to recognition of specific antigenic peptide on it by TCR. It should be noted that the hiding of the binding partner interface and utilization of the same interface to oligomerize and stand ‘upright’ on the cell surface, is a common strategy used by both CD4 (as described in the previous section) and pMHCII. This way, the superdimer of pMHCII stabilizes the ‘upright’ orientation for an efficient presentation of the antigenic peptide to T-cell. Therefore, the repeated observation of superdimers of pMHCII appears to be physiologically relevant, however, the presumption that pMHCII superdimer is the signaling unit led to models that did not explain subsequent findings. Similarly, in the case of CD4, the investigations supporting both D4-D4 pairs as well as D1-D1 pair appears to be correct and together with several structures, they appear to represent the physiologically relevant pre-liganded association.

Another important observation that emerged out of this analysis is that, in the ternary complex of CD4-pMHCII-TCR, the overall structure of monomeric CD4 (D1-D4 domains) is very similar to the apo structure of CD4 (RMSD ~1.8Å) **(Fig. S3-A)**[54]. Our analysis indicates that the D4 domain of CD4 which appears to interact with the membrane along its elongated surface as described in the apo structure, upon interaction with pMHCII moves away from the membrane, orienting itself almost perpendicular to the membrane surface **(Fig. S3-B)**. Although the overall structure of CD4 is unchanged, the reorientation of the D4 domain with respect to the membrane and the EJ-linker appears to provide sufficient length to reach the CD4 binding site on pMHCII present on the opposing cell. This reorientation due to interaction with pMHCII probably exerts sufficient machanoforce that dislodges the D4-D4 dimer consequently disturbing the relative orientation of its intracellular domains. This change in the intracellular part of CD4 might also help its association with Lck. Concomitant with this, it has been proposed in the crystal structure of the ternary complex of CD4-pMHCII-TCR, the arc-shaped structure also brings the C-terminal D4 domain of CD4 closer to the TCR:CD3 assembly.

### 1D array of CD2

CD2 is one of the first few structurally investigated cell surface molecules implicated in immune function and it is known to be present in close proximity to TCR:pMHC:CD4 complex on the cell surface [31, 60]. It is a cell adhesion molecule, expressed on T-lymphocytes. Human and rat CD2 are shown to interact with their cognate ligands CD58(LFA-3)[61] and CD48 respectively[62], all of which belong to CD2 family. The ectodomain of CD2 consists of a membrane distal IgV domain followed by membrane-proximal IgC domain. Overall structures of human (hCD2; PDB-entry: 1HNF) and rat CD2 (rCD2; PDB-Entry: 1HNG) are very similar **(Fig. S4)** with average C_α_ RMSD of 1.3Å, which confirms the relative rigid orientation of IgV and IgC domains [31, 60]. Interestingly, although CD2 is known to be a monomer in solution[63, 64], each molecule seen in both the structures is involved in the exactly similar dimeric association of IgV domains identical to that observed in canonical homophilic and heterophilic interactions involving IgV domains of immune receptors[65] and mimics the interaction of CD2 with its cognate partner CD58/ CD48 **(Fig. 4)**[31]. The superposition of rCD2 and hCD2 canonical dimers reveals that the two dimers are very similar **(Fig. S4).** The dimers can be depicted as either *trans* or *cis*, since the association satisfies both the forms **(Fig. S5-A) (See, supplementary information- section 4 for more discussion and examples on *cis*-*trans*notion)**. Based on the observation that the genes for CD58/CD48 and CD2 lie in proximity on chromosome-1[66, 67], it has been proposed that the heterophilic interaction in this family might have evolved from a common homophilic precursor[64], suggesting that the conservation of canonical homodimeric interaction observed in both CD2 structures, is probably physiologically relevant. Further, another structure of hCD2 containing only IgV domain maintains a similar canonical front-face to front-face interaction **(Fig. S4)**, even though the structure is domain-swapped (PDB entry: 1CDC)[68]. Maintenance of similar and canonical interactions between IgV domains in all the three different crystal structures suggest that the observed association, although very weak-to-be-detectable in the solution, appears to be the preferred mode of association of CD2 prior to its interaction with cognate partner.

**Fig.4.**
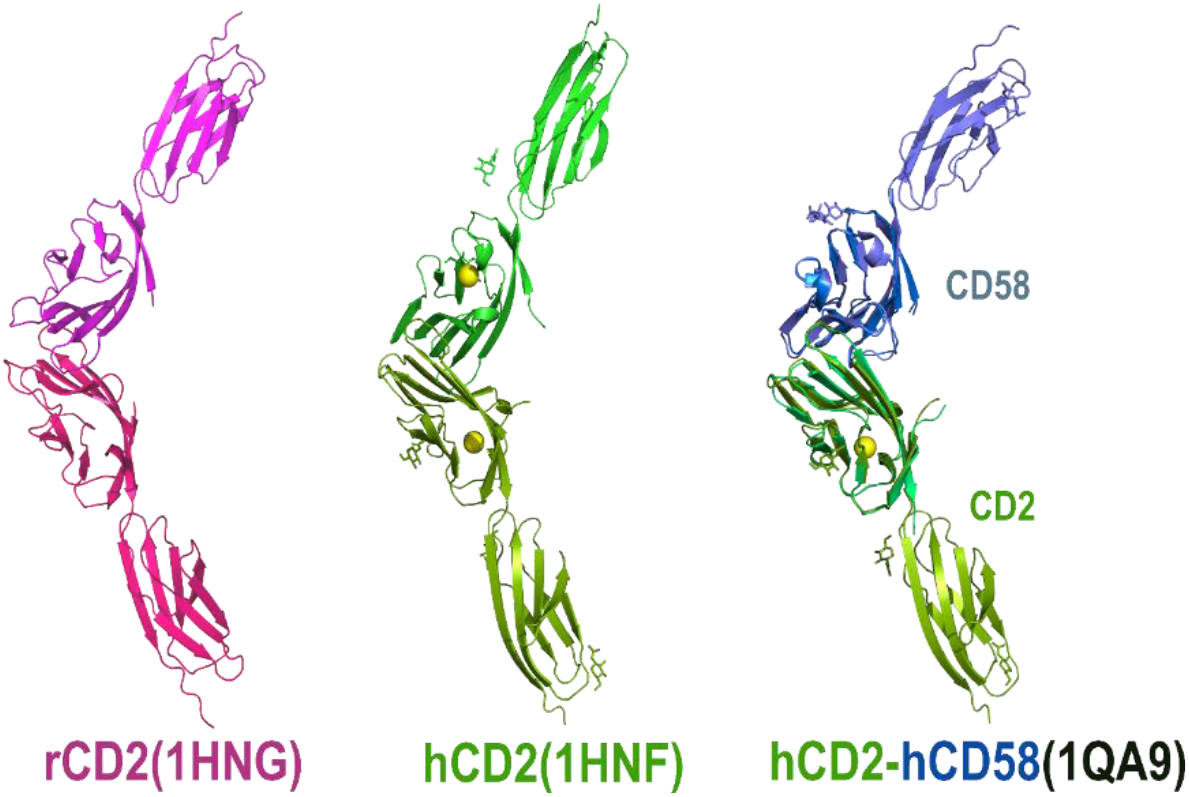
Comparative analysis of crystal structures of rCD2(PDB entry: 1HNG), hCD2 (PDB entry: 1HNF) and hCD2:CD58 complex (PDB entry: 1QA9). The dimeric associations observed in different crystals are similar and the homodimeric interactions of rCD2(magenta) and hCD2 (green) mimics the heterodimeric interaction of hCD2 with its cognate partner, CD58(blue) supporting the proposition that heterophilic interactions in CD2 family evolved from a homophilic precursor. hCD2:hCD58 structure shown here is obtained by superposition of full length hCD2 and full length CD58 on to respective proteins in CD2:CD58 complex structure containing only IgV domains (PDB entry: 1QA9)

Careful analysis of crystal packing of hCD2 molecules reveals that these molecules form a 1D array along the crystallographic *z*-axis, which places the membrane-proximal IgC domain (hence the EJ-linker) of all the molecules of the 1D array in the same direction as if they are emanating from the same cell **(Fig. 5)**. It is interesting to note that the N-glycosylation at Asn65(in the structure-1HNF) is located at the gap created between canonical CD2 dimers of this array and appears to act as a filler **(Fig. 5).** Conservation analysis reveals Asn65 is highly conserved (91%) across species **(Fig. 6).** In this array, the CD58 binding surface of CD2 is completely buried in the CD2 homodimeric interface. If the array seen in the crystal is reminiscent of pre-liganded state of CD2 molecules on the cell membrane prior to its interaction with CD58, then such an association with ‘closed’ ligand-binding surface might suffice the condition required for avoidance of spurious interactions and also provides support for ‘upright’ orientation of the molecules on the cell surface. Concomitant with this observation, single-molecule tracking of few raft-associated (Lck and LAT) and non-raft associated proteins (CD2 and CD45), which are known to be important in T-cell function, suggested an unexpectedly low diffusion coefficient for a non-raft associated protein CD2 (lowest among all the molecules studied)[69]. Here the assumption was that raft-associated protein should diffuse slowly compared to non-raft associated proteins. In this context, the 1D association appears to be physiological and provides an explanation for unusual low diffusion coefficient observed for the non-raft associated CD2.

**Fig.5.**
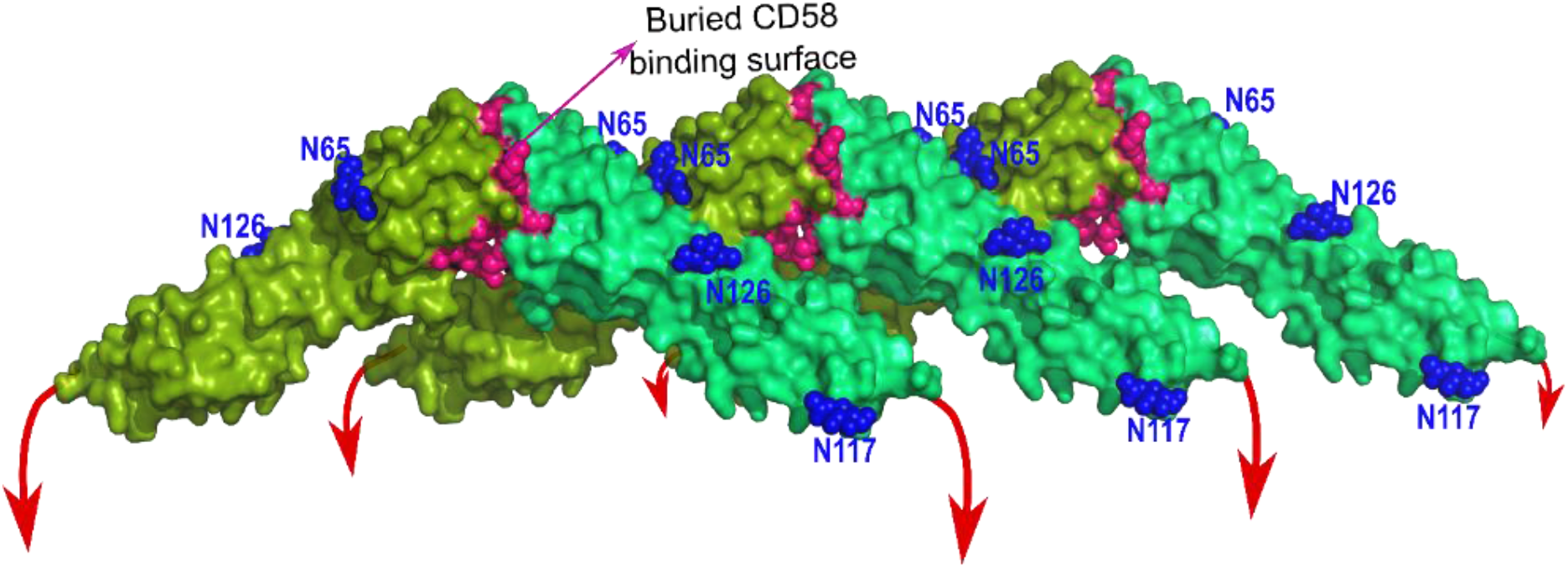
Surface representation of one-dimensional array of hCD2 as observed in the crystal structure (PDB entry:1HNF). Note that the dimer which appeared as ‘*trans’* as shown in Fig.4 can also be considered as ‘*cis’* and such *cis* dimers hide the CD58 binding site (shown as pink spheres) on CD2. The *‘cis’* dimers further interact to form a one-dimensional array of molecules supporting each other in an *‘*upright*’* orientation perpendicular to the membrane. The N linked glycans are shown in blue spheres. N-glycans (at N65) between the adjacent *‘cis’* dimers act as fillers and the glycan at N117 which is close to the membrane might stabilise the *‘*upright*’* orientation. The red arrows indicate EJ-linkers extending towards membrane.

**Fig.6.**
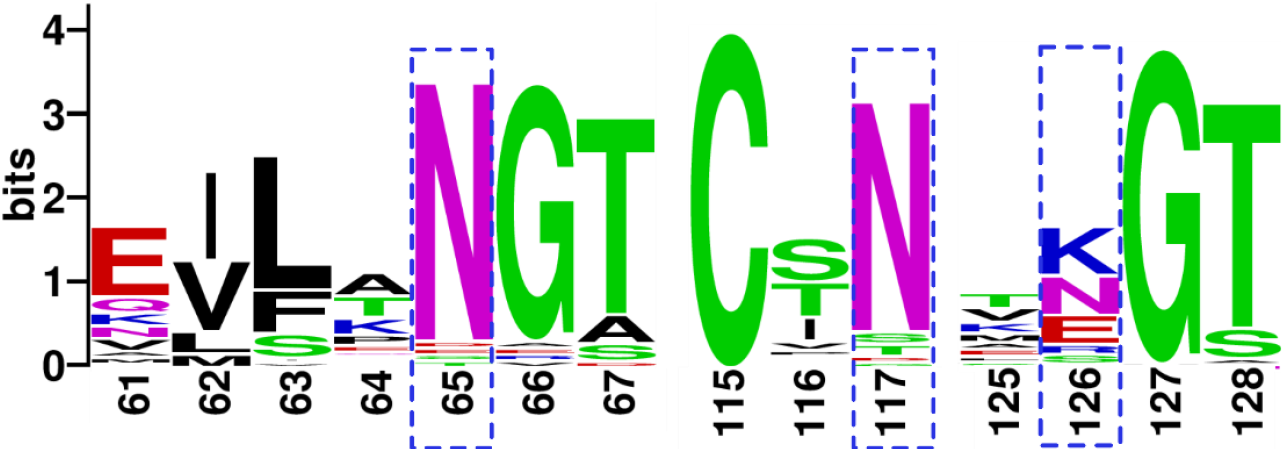
Sequence logo depicting the conservation of N-glycosylation sites in CD2 across species: N65 and N117 boxed in blue dashed lines (numbering according to PDB structure 1HNF) are conserved by 91% and 86.4% respectively. While the glycan at N65 might act as a filler between the adjacent proteins of the 1D-array, the glycan at N117 lies close to the membrane and might have stabilising interactions with the membrane. The third glycan is at N126, also seen in the structure, but the site is only 27% conserved.

### 1D array of TNFR1

Tumour Necrosis Factor (TNF) is a cytokine which interacts with its receptors TNFR1 and TNFR2. Currently, several TNF family members (29 receptors and 19 ligands in humans) have been identified and are known to play an important role in the immune response. While the ligands are type-II membrane proteins with trimeric extracellular domain and include both membrane-tethered as well as soluble versions, the receptors are type-I membrane proteins mostly membrane-bound with the exception of soluble decoy receptors. Also, receptors are elongated structures with 3-4 cysteine-rich domains (CRDs) and some of them interact with multiple ligands[70]. Further, the ‘ligand-induced trimerization’ model suggests that the receptors get trimerized only in the presence of trimeric ligand, where the inter-digitization of monomeric receptors in the groves of trimeric ligands have been observed[71]. It has been proposed that the common trimeric architecture of TNF family ligands together with the promiscuous binding of receptors to different ligands creates the possibility of the formation of deleterious mixed trimers [72]. TNF mediated signaling being involved in a variety of processes ranging from cell proliferation to cell death induction, the formation of deleterious mixed-trimers put forth a challenge that how the receptors avoid the formation of mixed trimers to generate robust signaling? This raises the question of whether there exists any pre-liganded assembly of TNFR1 that can segregate specific TNF receptors prior to the ligand-binding and avoid mixed trimeric assembly of ligand and receptors?

The crystal structure of TNFR1(PDB-1NCF)[73], reveals parallel dimeric association with predominant interactions located on the N-terminal CRD1 **(Fig. 7A).** Consistent with this observation, based on the mutational studies on the CRD1 and based on complete replacement/exchange of this domain with the corresponding domain from the related receptor, it has been proposed that CRD1 is responsible for ligand-independent assembly of the receptor, and hence named as ‘pre-ligand binding assembly domain (PLAD)’[74]. In brief, what is evident from various studies is that in the TNF family, the control of the signaling event is overly complex and is not mediated by mere ligand-receptor interactions. This further supports the possibility of the existence of a pre-signaling assembly of specific TNF receptors, prior to the ligand-binding and which is distinct from the ligand-induced trimer. The PLAD mediated cheek-to-cheek association of TNFR1 appears to satisfy the condition required for stabilization of the ‘upright’ orientation of this receptor on the cell surface. However, the existence of a long EJ-linker in TNFR1 (15 residues) raises the question that how such a long linker stabilizes the ‘upright’ parallel orientation of TNFR1 on the cell surface (TNFR2 has 56 residue EJ-linker)?

**Fig.7.**
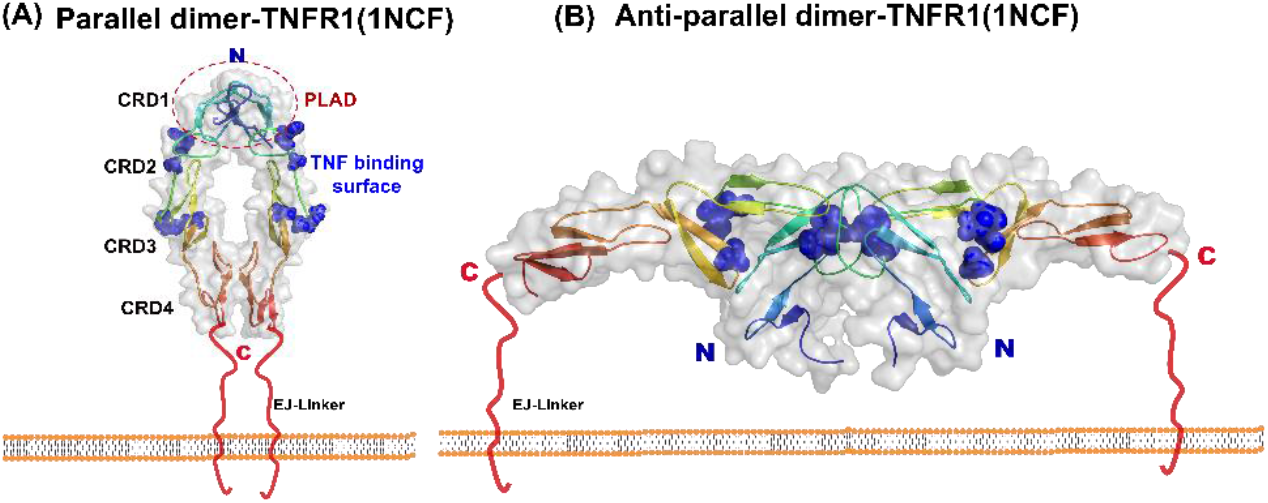
Different dimeric interactions observed in the crystal structure of TNFR1. **(A)** Parallel dimeric association of TNFR1 (PDB entry: 1NCF) interacting via the N-terminal PLAD (encircled in red dashed lines) exposes the ligand binding surface (residues shown as blue spheres). **(B)** The anti-parallel association, also observed in the same crystal structure effectively hides the ligand binding residues in the dimeric interface.

From the TNFR1 crystal structure, Naismith *et. al*, also recognize an association that can be assigned as an anti-parallel association of the receptors, which buries the ligand biding surface distributed over both CRD2 and CRD3 domains **(Fig. 7B).** Based on this observation, they propose that the burial of ligand-binding residues might avoid unintended signaling events[73]. The anti-parallel model is also convincing as it explained the differential cross-linking effects of therapeutic agonistic and antagonistic antibodies[75]. However, the anti-parallel model undermines the role of PLAD which was proven to be physiologically important by many experimental studies. Our packing analysis reveals an interesting observation, where PLAD mediated parallel-dimers associate with each other burying the ligand-binding site and forms supramolecular 1D array containing both parallel and anti-parallel dimers **(Fig. 8).** Such an ordered and continuous association of molecules, together with the long EJ-linker might help these molecules to stand ‘upright’ and segregate on the cell surface. Further, the zipper like arrangement with hidden active sites might only respond to the presence of sufficient concentration of specific TNF ligands that probably unzips the 1D array. Again, this observation reconciles two contradicting experimental observations that (1) PLAD mediated parallel dimerization is important for function and (2) anti-parallel association is physiologically relevant. Further, the 1D zipper mediated segregation of specific kinds of TNFRs might also avoid the deleterious mixed-trimer formation of ligands and receptors. Supporting this observation, another crystal structure of TNFR family member called DCR3 maintains both parallel and anti-parallel dimeric association (PDB-3MHD) **(Fig S8)**, probably suggesting a common association preference of TNFR family members. The role of transmembrane domain of the TNF receptors in stabilisation of ligand induced trimeric association is discussed in the **supplementary information- section 5.** Hence, signatures at the TM domain that support the trimerization and those at the ectodomain that supports dimerization, being facile, appear to facilitate the required oligomeric organization depending on the pre- and post signaling states.

**Fig.8.**
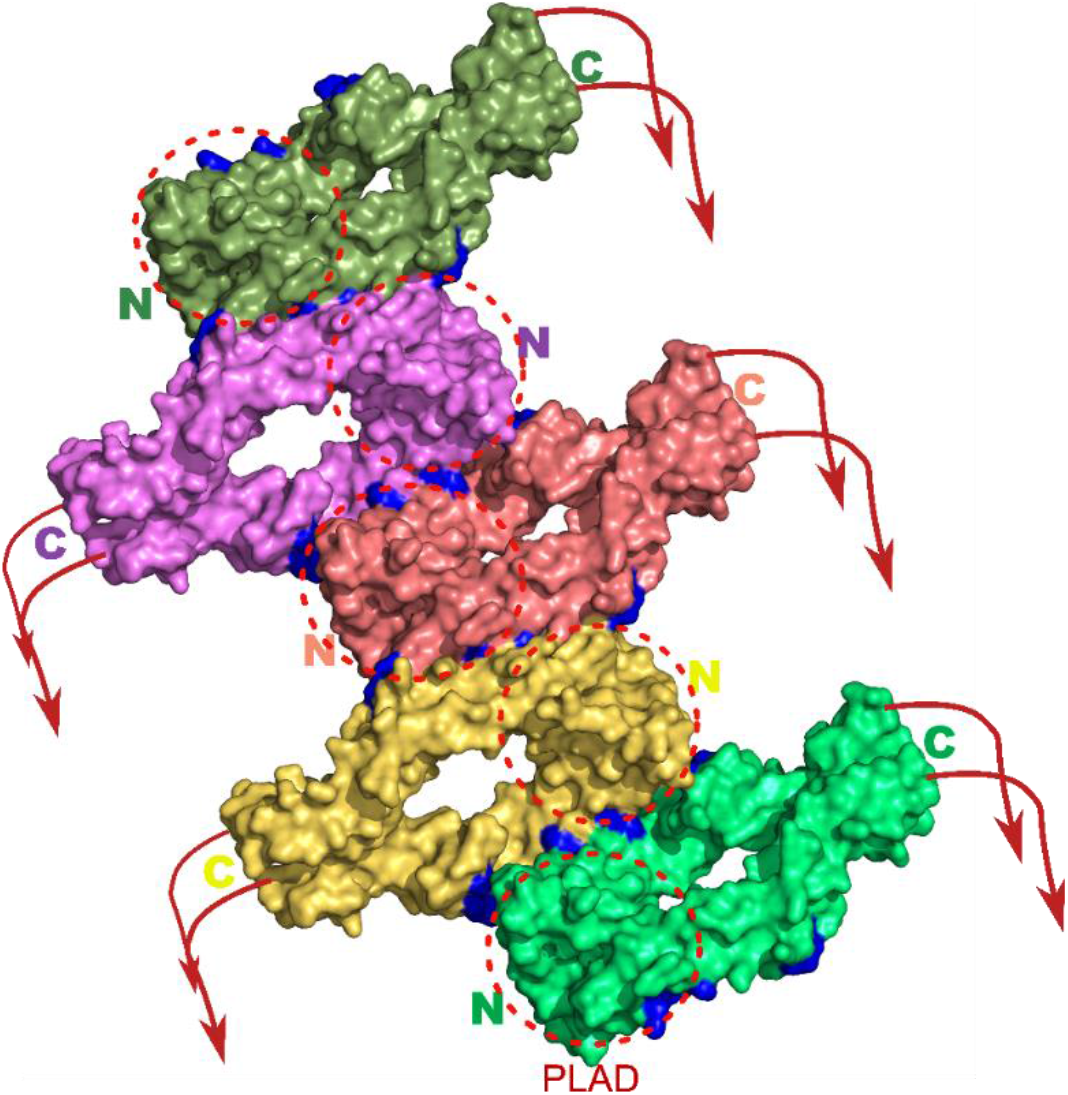
One dimensional array observed in the crystal structure of TNFR1 that supports both parallel and antiparallel dimeric associations. Such an array of molecules on the cell membrane stabilized by velcro of weak interactions support the upright orientation of receptors and accounts for the importance of PLAD (encircled in red dashes) in receptor association as well as hides the ligand binding site (spheres in blue) to prevent spurious signalling and thus reconciles the opposing views.

Recently, various inhibitors are designed to manipulate the PLAD assembly and have been shown to inhibit downstream signaling [76][77][78], suggesting the existence of PLAD assembly on the cell membrane. Further, hereditary heterozygous mutations in the Fas receptor, which also belongs to the TNFR1 family (explained in the main text), exhibit a dominant-negative effect leading to Autoimmune lymphoproliferative syndrome (ALPS). In patients with ALPS disease, the mutant Fas receptors cannot bind to the Fas ligand. But, since they are heterozygous mutants, when co-expressed along with wild type(wt) receptors, the non- ligand binding mutant Fas receptors were found to interfere in the apoptosis activity of wt receptors indicating that the mutant receptors can interact with the wt receptors even prior to the ligand-induced trimerization[72]. These observations indicate the *cis* interaction of the Fas receptors on the cell surface.

Apart from the specific examples described above, the mode of association of several cell surface receptors observed in the crystal structures mimic the condition required for stable ‘upright’ orientation. One of the classic examples that clearly depicts the importance of crystallographically observed *cis-*association of TIGIT that was later proven to be important for signalling is also discussed. **(supplementary information - section 6 and Fig. S6)**.

Our analysis also suggests that stabilization of ‘upright’ orientation can be mediated by any part of the ectodomain, and appears to be optimally carved for its size, architecture, and functional role. Adding to these possibilities, small molecules and ions in the extracellular milieu can also contribute to the specific organization of cell surface proteins. Similarly, the negative charge on glycans is expected to repel from the membrane and probably help the weakly associated molecules to stand ‘upright’[79] **(See, supplementary information- section 7&8)**

## Conclusion

Spontaneous organization is a general phenomenon observed at all length scales [80, 81]and is fundamental to all living cells, where their shapes, motility and functions are controlled by protein assemblies. Even the cell surface is compartmentalized to exert versatile and robust function. Here, we observe that the cell-surface molecules oligomerize to support appropriate positioning of the membrane distal domains for robust signaling, effectively avoiding spurious signaling events. More interestingly, the *cis-*association of these molecules coherently reconcile contradicting observations suggesting that the oligomerization and often formation of an ordered array is a common phenomenon, probably utilized by many (if not all) of the cell surface molecules involved in signaling. However, none of these proteins show such oligomerization propensity in solution, suggesting that the observed *cis-*interactions are too weak-to-be-detected by biophysical studies. This weak oligomerization propensity appears to be essential as these molecules are required to organize/reorganize depending on the signaling state. Supporting this observation, as discussed earlier, the mutation studies to understand these interactions as well as those mutations associated with pathological conditions resulted in impaired signaling. The analysis and the conclusions drawn here entirely relies on the repeatability of a pattern, i.e., the pattern of association in several crystal structures of the same or related molecules, and repeated observation that constitutively expressed cell-surface molecules oligomerize and often bury their ligand-binding surface and so on. To retrieve a pattern and matching them requires sufficient structural and functional data on these molecules to efficiently connect the results of the pattern of association to their functional roles. For example, pMHCII and TCR are molecules involved in the continuous screening process; hence, the self-association of pMHCII into higher-ordered arrays may not support efficient peptide presentation. Similarly, the pMHCII recognition site on TCR should always be accessible. Accordingly, neither the pMHCII superdimer nor the TCR gets involved in a supramolecular organization. Since, the antigenic peptide already provides required specificity between TCR and pMHCII, an additional restriction may not be required. In contrast to specific TCR:pMHCII interaction, the binding of CD4:pMHCII is non-specific. Accordingly, CD4 function is regulated by its *cis*-oligomerization as well as by the formation of pre-liganded 1D array, which hides its pMHCII biding surface in the non-signaling resting state. Hence, CD4 interacts with pMHCII only upon proper recognition of antigenic peptide by TCR that opens the CD4 binding site on pMHCII. In the case of TNFR1, a constitutively expressed protein, the observed 1D array suffice the requirements for both the contradicting observations as well avoids spurious signaling. Hence, it appears that these signaling molecules are carved with well-defined surface chemistry and complementarity, the strength of which appears to be optimal for their functional role. Therefore, what is generally termed as cluster of molecules on the cell surface, without any impetus to their specific order, appears to be inappropriate and models such as ‘lipid-rafts’ and even the ‘fence and pickets’ may not provide an explanation that emulates specific sorting and robust signaling ability of these molecules. Although thousands of experimental work supporting these models co-exist with unhealthy controversies[14], in the absence of know-how on their ordered association preferences, more experiments and controversies continue to spring. It can be arguable that the observed effect of ‘lipid-rafts’ or ‘fences and pickets’ or ‘molecular crowding’ is probably a consequence of the inherent ordered association of cell surface proteins that places their relatively rigid transmembrane domain in regular and ordered fashion to which molecules such as sphingolipids and cholesterol can adhere to.

Similarly, as observed in the crystal structures of hCD2, cadherins, ephrin and so on, when these molecules are free in solution, at higher concentrations they can form large arrays of the dimension of the crystal itself (~100 microns). In contrast, when they are tethered to a membrane, the dynamics of cytoskeleton supported active membrane might limit the growth of these weakly associated arrays or due to the curved membrane architecture. Hence, it appears that ‘fences and pickets’ imagined out of cytoskeleton elements, probably can control the growth of these clusters, but may not carry an element of their organization specificity and functional control. Consonant with this observation, recent studies demonstrate that when the cytoskeleton is disturbed by cytoskeleton inhibitor ‘latrunculin A’, cadherin clusters were found to grow to a size that is larger than generally observed [82].

Overall, it appears that the clustering being the key element of signal processing; the rules that govern the clustering, their type, optimum strength, accuracy and reproducibility of the signal are hidden within the defined set of self-organizations of cell surface molecules. Such an organization is maintained by surface complementarity of the molecules together with polar interactions that are facile enough to allow their organization/reorganization depending on the functional context. The role of much neglected EJ-linker in *cis*-association of these molecules is also described here, suggesting elements of weak association can be located anywhere on the intracellular, TM and extracellular part of the molecule or cumulative effect of all these interactions. The presence of cognate-partner with higher affinity and appropriate concentration might dislodge the pre-liganded *cis-*interactions and reorganize into an array of different architecture, strength and dimension and consequently position their intracellular cytoplasmic domains in an ordered fashion, the altered geometry of which might support the recruitment of downstream signaling molecules. Considering these observations, we would like to call our model as “Preferential Protein Interaction of Signaling Receptors” (**PPISR-Model**). In the absence of a suitable model that can explain the cell surface heterogeneity and its importance in function, we believe that this new conceptual framework solely developed based on the available data and recurrence of pattern, appears to provide a basis for future experiments for molecules not only involved in immune function but also for a repertoire of other signaling receptors. Further, the knowledge of the physiological association of the cell surface receptors in their resting state is very important to understand the signalling mechanism and also in the light of drug design. Mostly, the inhibitory antibodies are designed for blocking the active sites of the receptor to preclude its binding to the cognate partner. If the observed *cis-*association in which the binding site is buried is true, then the inclusion of this knowledge in drug design will be more effective. This is exemplified in the case of TNFR2 receptors by the requirement of at least a F(ab)2 antibody rather than a single Fab fragment to effectively lock the physiological confirmation and render antagonism[75, 83]. Further, such an understanding will form the basis for the design of drugs that disturb or stabilize the weak pre-liganded assembly.

## Abbreviations

IgV: Immunoglobulin V-type domain
IgC: Immunoglobulin C-type domain

## Acknowledgment

UAR would like to thank DBT, India for Ramalingaswamy fellowship (2011-2015) and Vision Group on Science and Technology (VGST), grant #191, Karnataka, India. SL would like to acknowledge the fellowship from CSIR, Government of India. The authors would like to acknowledge Admar Mutt Education Foundation (AMEF) for the facility and support.

## Author contribution

UAR conceptualized and reviewed the manuscript. UAR and SL analysed the data and wrote the manuscript

## Supplementary Materials

Methodology

Figures S1-S10

