## supplementary material for "How does an ectodomain of membrane-associated proteins stand upright and exert robust signal?"

**Author Information**

**Swetha Lankipalli[1,2] and Udupi A. Ramagopal[1 *]**

[1]Biological Sciences Division, Poornaprajna Institute of Scientific Research (PPISR), Bidalur post, Devanahalli, Bengaluru, 562164, India

[2]Manipal Academy of Higher Education, Manipal, Karnataka, India-576104

*To whom correspondence may be addressed:

Udupi A. Ramagopal, Biological Sciences Division, Poornaprajna Institute of Scientific Research (PPISR), Bidalur post, Devanahalli, Bengaluru, 562164, India
Ph.: +91 9900810182;

**Abbreviations:**

IgV- Immunoglobulin V-type domain; IgC- Immunoglobulin C-type domain

**Keywords:** Ectodomain; ‘upright’ orientation, protein clustering; one-dimensional array; nanoclusters, spurious signaling; non-signaling resting state.

### 1. Methodology:

**1.1 Structural analysis of cell surface molecules**: To understand the association preference of the cell surface receptors, various structures of these molecules were downloaded from the Protein Data Bank (wwPDB)(*1*) and their packing preferences were analysed using the program COOT(*2*). Similarly, the packing preferences of non-membrane associated proteins were also looked at to know, if any, preferential orientation of cell surface receptors can be observed. To select proteins that are known to be monomers in solution, we reviewed the literature to verify whether the monomeric nature of these proteins in solution was supported by different biophysical and biochemical experiments. Further, only those proteins for which multiple crystal structures are available, either from the same organism or from closely related ones are considered for further investigation. The structure of molecules containing all the ectodomains and those containing only few domains were separated. None of these structures contained the extracellular juxtamembrane linker (EJ-Linker). The structures of the molecules containing all the domains of the ectodomain were compared with the truncated ones using SUPERPOSE program of CCP4i suite(*3*) and also with the structure modelling program COOT, to understand the similarity of the structures. Similarly, the complete ectodomain from different crystal structures were also compared to understand the similarity of these structures crystallized in different conditions and space groups using the above programs.

**1.2 Sequence conservation analysis:** To analyse the conservation of glycosylation sites in hCD2, the corresponding ectodomain sequence with uniport ID: P06729(amino acid range: 25-209) was given as the query in BLAST(*4*) and 5000 hits were generated using non-redundant database. The resultant initial 5000 hits were further filtered to obtain sequences with a percent identity cut-off of min 50% - max 99.5% and with a query coverage of minimum 80%. The resultant sequences were used in CD-Hit(*5*) to generate clusters with sequence identity cut-off of 90%. The representative sequences of the clusters generated from CD-Hit were aligned using Clustal Omega(*6*) and analysed using Geneious(*7*). Sequence logo was generated using WebLogo(*8*).

**1.3 EJ-linker sequence analysis:** Since the boundaries of EJ-linker were not defined clearly in UNIPROT, we deduced the sequence length from the available details. The region between the sequence mentioned as transmembrane domain and the extracellular domain was considered as EJ-linker and cross-checked using available crystal structures. From the resulting sequences the percent of prolines and hydrophobic amino acids in the sequence was calculated.

### 2: Literature, current advances and unsolved mysteries: A search for new model and a reason for analysis of association seen in crystal structures.

**2.1 Super-resolution microscopy reveals diverse nano, meso and microclusters of cell surface molecules:** Advent of new techniques, sophistication of instrumentation and development of better theoretical approaches have revolutionized the fields such as super-resolution microscopy, single molecule tracking and so on. As the methods to look at these molecules are improving; more diversity and complexity of the protein clusters are getting revealed. For example, cadherin clusters once thought to be initiated by trans (cell to cell) interactions, followed by diffusion trap leading to the formation of trans/cis microclusters, now appears to be different in that the larger *trans-cis* microclusters co-exist with lateral (*cis*) nano-clusters (*9*). Similarly, in the case of endothelial receptor CD36, its interaction with multivalent ligands, such as TSP-1 (thrombospondin-1) was shown to result in clustering and consequent downstream signaling (*10*). However, recent super-resolution microscopy studies reveal that CD36 nanoclusters are enriched with clusters of downstream effector molecules 'Fyn' but only the larger and denser clusters of CD36 induced by TSP-1 binding activates Fyn(*11*). A similar observation on TCR-CD3 complexes, where only denser clusters were found to be phosphorylated and associated with downstream signaling was reported(*12*). Similarly, in the case of β2-integrin LFA-1(Lymphocyte function-associated antigen-1), three different avidity patterns: randomly distributed inactive molecules, ligand-independent ordered proactive nanoclusters and the macroclusters triggered by ligand interaction(*13*) have been observed. These observations suggest probable pre-liganded assembly of these molecules on the cell surface which is distinct from post-liganded assembly.

#### 2.2 Examples of cases where crystal packing revealed physiological interactions:

Crystal packing sometimes reveals physiological interactions. In many ectodomain structures of homophilic immunoglobulin superfamily cell adhesion molecules, the molecules in the crystal arrange in a zipper-like array representing their functional role in the cell-cell junction(*14*). Similarly, crystal structures of CTLA-4 in complex with B7-1 and with B7-2, two functionally equivalent molecules, revealed identical 1D array consisting of alternative dimers of B7s and CTLA-4(*15*, *16*), representing the possible arrangements of these molecules during the immunological synapse formation. In all these cases, the *trans* (between the molecules from opposing cells) interactions are strong enough to be detected in solution. Hence, it can be inferred that the relatively strong receptor: ligand (homophilic or heterophilic) interactions can survive and, in most cases, override the crystal packing forces, representing the expected association *in vivo*. Although the *cis* interactions are too weak-to-be-detected in solution, the high concentration of proteins on the cell membrane, presence of other proteins (excluded volume effect) and the restricted orientation of proteins due to membrane tethering increases the likelihood of their oligomerisation by several orders (*17*). Hence, we analysed the crystal structures for the presence of any preferred interactions which might represent the pre-liganded physiological *cis* associations.

**3: The ectodomain of a cell surface receptor is not always flexible as presumed.**

**3.1 Few examples of proteins with rigid inter-domain interactions:** Analysis of two different crystal structures of Nectin-1 ectodomain (D1-D3), with the first structure containing one molecule in ASU (PDB-entry:3U83)(*18*) and the other with four molecules in ASU( PDB-entry: 4FMF)(*19*), revealed rigid interdomain interactions. The superposition of complete ectodomain of Nectin-1 in two different crystal forms results in $C_\alpha$ RMSD of less than 2.0Å. It was also interesting to note that the structure of D1-D3 domains of Nectin-3 (PDB-entry: 4FOM) is very similar to Nectin-1. In the case of CD22 with seven Ig domains, although the structure of complete ectodomain is not available, the apo (PDB entry: 5VKJ) and sialyllactose bound structures (PDB entry: 5VKM) having D1-D3 domains are very similar. In line with this observation, using small-angle scattering studies and negative stain electron microscopy; it has been shown that full-length CD22 ectodomain (D1-D7) adopts similar rod-like conformation with limited interdomain flexibility (*20*).

**3.2 Examples illustrating importance of EJ-linker:** The possible role of EJ-linker can be observed in, for example, B7-1 and B7-2 which are functionally equivalent molecules. Upon interaction with CTLA-4, both B7-1 and B7-2 form a similar kind of 1D zipper-like organization consisting of an alternative arrangement of B7 dimer and CTLA-4 dimer mimicking the situation expected for these molecules in the immunological synapse(*15, 16*). However, while B7-1 is a known dimer, B7-2 is shown to be a monomer in solution (*21, 22*). Interestingly, the human B7-2 linker region ($_{237}$EDPQPPPDHIP$_{247}$) contains 54.54% hydrophobic residues (including proline) and 45% prolines. Proline is an anomalous residue (average occurrence of around 6%(*23*)), and proline-rich sequences such as collagen, keratin and so on are known to associate in parallel.

Unfortunately, constructs of B7-2 used in the solution experiments to understand the B7-2 oligomerization did not have this proline-rich sequence. Even in the FRET-based studies(*24*), aimed to reveal the oligomerization of B7-2 on the cell surface, the EJ-linker sequence has been modified to a sequence rich in -Gly-Ala-, thus leaving no clue on the effect of original EJ-linker on B7-2 oligomerization on the cell membrane. Similarly, the EJ-linker of TNFR2 contains 56 residues with 50% hydrophobic residues of which 12 are prolines (~22% of EJ-linker). A recent study that provides a model for TNF superfamily signaling based on the structures of different TNF receptors does not consider the importance of the probable structure of such a long linker. However, the importance of the EJ-linker (also termed as stalk) in the case of TNFR-1 and TNFR-2 proteins has been demonstrated in a study where the exchange of the stalk regions led to the change in their responsiveness to the membrane-bound ligand(*25*).

**4. *Cis* and *trans* convention:**

The description of interactions observed in crystal structures as *cis* or *trans* in case of homophilic molecules appears to be inappropriate as the same interaction can be depicted as *cis* or *trans*. While rCD2 dimer mimicked a *trans* interaction with its cognate partner CD48, the highly similar hCD2 dimeric association represented *cis* interaction of the molecules on the cell surface **(Fig. S5-A).** This observation suggests that differentiating these interactions as either *cis* or *trans* appears to be inappropriate and sometimes leads to wrong interpretations. For example, in the structure of the D1-D2 fragment of CD4, it was concluded that the observed interaction between the D1 pair is *trans* as they appeared to come from two different cell surfaces (D2 domains are placed in opposite directions). However, this structure superposes well with the *cis* dimer observed in the crystal structure of complete ectodomain (D1-D4, **Fig. S2-A**) of CD4(*26*). Here, we quote few examples of interactions which can be visualised as both *cis* and *trans* but have been generally labelled as *trans*. NTB-A also called SLAMF6, is a homodimer and member of the CD2 family. In the crystal structure, the angle between the two protomers of the dimer is approximately 86°. The homomeric interaction seen in the crystal is always represented as *trans* interaction as if two molecules are coming from juxtaposed cells **(Fig. S5-B)**(*27*, *28*), However, it is also possible that they can stand on the same cell surface as dimers using the same canonical dimeric interface as observed for human CD2. Hence, it is likely that the homomers can sometimes use their membrane distal domains for both *cis* and *trans* interactions, and in the absence of *trans* interaction with juxtaposed cells, they might utilize the same surface for *cis* interaction and stabilization of ‘upright’ orientation and avoid spurious signaling. Structures of homophilic dimers such as Nectin-1, CD166 are shown in **Fig. S6-A, S6-B**, representing both the possible orientations on the cell membrane.

**5.0 Transmembrane domain mediated interactions of TNF receptor superfamily members:**

Although there is considerable debate on whether TNF receptors exists as parallel dimers or anti-parallel dimers (which we think is resolved with our model) in the pre-signalling resting state, the question that, how these dimeric receptors involve themselves in 'ligand-induced trimerization' remains unclear. Various studies on single-pass transmembrane (TM) domains have shown that their organization does play an important role in the function of these proteins. (*29-31*). In case of CD95 (FasR), an apoptosis-inducing death receptor and a member of tumor necrosis factor receptor (TNFR) superfamily, the NMR structures of both human and mouse TM domain revealed a highly analogous homo-trimeric organization(*32*). Structure-guided and cancer-related mutations that disturb the TM trimerization and *in vivo* studies demonstrated that these mutants show impaired apoptosis induction, suggesting the importance of trimerization of TM domains in signaling (*33*). Sequence analysis of different TNFR superfamily members reveals a conserved proline in each family at a specific position on the TM domain, indicating that proline plays a critical role in the trimerization of TNFR superfamily members. Based on the extensive analysis, the authors conclude that the TM-trimerization helps in post-ligand trimeric associations of FasR with its ligands including Fas, which are again trimers, resulting in clustering of liganded FasR:FasL complexes(*32*). They also propose that the pre-liganded upright dimer of the ectodomain, re-associates into trimer in the presence of Fas ligand. Hence, signatures at the TM domain that support the trimerization and those at the ectodomain that supports dimerization, being facile, appear to facilitate the required oligomeric organization depending on the pre- and post signaling states. Consonant with these observations, super-resolution microscopy data shows that the dimers and trimers of FasR co-exist on the plasma membrane(*31*).

**6.0 Other examples illustrating *cis* interaction mediated 'upright' orientation:** In addition to the examples like CD4, pMHCII, CD2 and TNFR1 explained in detail in the main text, there are many other receptors in which such *cis* interactions are shown to mediate the 'upright' orientation. For example, the extracellular domain of T cell immunoglobulin and ITIM domain (TIGIT) contains a single IgV domain, with a 15 residue EJ-linker. It is an inhibitory receptor expressed on the surface of natural killer cells (TIGIT:CD226/Nectin-2:PVR pathway is analogous to CTLA-4:CD28/B71-B7 pathway, where both CTLA-4 and TIGIT are inhibitory receptors on T-cells). TIGIT is known to be a weak homodimer, which is thought to dimerize using canonical front-face to front-face (*trans)* interaction observed in IgV domains (based on SPR and analytical ultracentrifugation studies). There are three crystal structures of human TIGIT crystallized in three different conditions and space groups. Among these three cases, (PDB-entry: 3UCR, 3Q0H, and 3RQ3), two of which even miss the front-face to front-face interaction described for this class of molecules (also described as "lock-and-key" mode of interactions). While the expected front-face to front-face interactions in three different crystal structures of TIGIT are different, TIGIT:TIGIT *cis*-dimer is observed in all three structures mentioned above and in the structures of TIGIT:PVR(PDB-3UDW) and also in the TIGIT:Nectin-2 structure(PDB-5V52) (total of 5 structures, Fig. S7). That is the back-face to back-face association (*cis* homodimer) is conserved in all five structures. The structure-guided mutation on TIGIT based on the TIGIT:PVR structure (I42A and I42D mutants on TIGIT) **(Fig. S7)** that disrupt the TIGIT *cis*-dimer is shown to limit TIGIT:PVR mediated cell-adhesion and TIGIT induced PVR phosphorylation in primary dendritic cells, although it binds to PVR with similar affinity (TIGIT interacts with PVR through its front face). Further, while clustering was observed at the junction of cells expressing wtTIGIT and PVR, reduced clustering was observed with both I42A and I42D mutants. The observation of cis-dimer in 5 different crystals structures and cell-based experiments together confirm that the weak-dimer observed in solution studies is probably a *cis* dimer, disruption of which results in impaired signaling and clustering. This is a classic example where *cis*-dimer disrupting mutants generated purely based on crystallographic observation have shown to impact their organization on the cell-surface(*34*)

In a similar example, there are four structures of PD-L1 containing both IgV and IgC domains crystallized in different conditions (PDB entries: 4Z18, 3FN3, 5JDR, 3BIS)(*35-37*). It is interesting to note that in all the four cases, PD-L1 crystallizes as a parallel dimer (*cis* dimer), where the membrane-proximal C-terminal ends are close to each other **(Fig. S9-A).** Although no biophysical experiments can detect their dimeric association in solution, crosslinking studies show mostly the dimeric population(*35*). However, the possible reason for such an association has never been described before. Our analysis of the structures reveals that the back-face of the IgC domain consisting of ABED strands from both the interacting molecules fuse together to form a continuous β-sheet stabilizing the expected physiologically relevant dimeric structure that can support 'upright' orientation **(Fig. S9-B).** The similar, but slightly distinct fusing of ABED strands from the IgC domain has also been observed in the structure of the full-length ectodomain of CD58. Similarly, in the crystal structures of human EphA2 receptor (Eph, a cell surface receptor protein tyrosine kinase) both the liganded and unliganded forms show *cis* contacts between ligand-binding domains (LBD- LBD), sushi domains (sushi-sushi) of adjacent proteins. Similar to the perching of the D4 domain of CD4 on to membrane, in the case of the EphA2 receptor, the membrane-proximal FN2 domain is oriented almost perpendicular to the other four domains of the structure(*38*). In a recent study, using MD simulations and mutational studies it was shown that the FN2 domain interacts with membrane along its elongated surface through positively charged lysine and arginine residues and thus stabilizes the Ephrin dimers on the cell surface **(Fig. S10)**(*39*). It can be noticed that as the size of the ectodomain increases (four domains in CD4 and five domains in Ephrin), other than the oligomeric association, nature sometimes appears to utilize direct interaction of membrane-proximal domain with the plasma membrane as an additional stabilization factor. In the case of CEACAM1, SPR based binding studies, chemical cross-linking experiments and molecular electron tomography studies on CEACAM1 confirms the adaptability of ectodomain to make parallel *cis* interactions (A-dimers) as well as anti-parallel *trans* dimers (C-dimers). Liposomes with anchored CEACAM-1 molecules also have shown clusters and the degree of clustering did not change with change in lipid quantity suggesting that the clustering is influenced more by ectodomain interactions rather than by protein-lipid ratio on the membrane(*40*).

### 7.0 Metal ions play important role in structural stabilization of ectodomains and clustering.

The role of metal ions in stabilising the *cis* mediated interactions is demonstrated in the case of many cell surface receptors. For example, in the case of cadherins, interdomain stabilization is mediated by calcium ions(*41*) and hence most crystallization soup, and consequently, the crystal structures are derived in the presence of $Ca^{2+}$ ion. From these structures, it is clear that $Ca^{2+}$ ions play a critical role in interdomain stabilization. Also, it has been observed that the presence of cations, such as $Ca^{2+}/Mg^{2+}$ contributes to the stability of CEACAM-1 ectodomains and increases the formation of dimers and multimers(*40*). The role of $Ca^{2+}$ ions in the organization of membrane-associated proteins like Syntaxin and SNAP25 have been reported and its role in higher-order clustering has also been discussed(*42*). This situation calls for the use of physiological concentration of $Ca^{2+}$ in other experiments such as super resolution microscopy to effectively compare the results from independent experiments.

### 8.0 Role of glycans in stabilising upright orientation:

Considering the hydrophilicity, flexibility, mobility of glycans in the aqueous environments, the glycans are expected to be the best fillers that can allow adaptation to changing environments, depending on the organization/reorganization of these receptors on the cell surface. For example, in the case of symmetric dimers, such as CD2 and NTB-A any glycan attached to the membrane-proximal domain of one of the protomers of the dimer places the glycan attached to another protomer in the opposite direction, creating similar glycan-membrane interactions at the two opposite sides of the plane passing through both the protomers of the dimer and perpendicular to the membrane **(Fig. 5 main text)**. Hence, glycans might also play a role in the erect orientation of the molecules on the surface, however, considering their flexibility, they may not by themselves define the preferential association.

### Supplemetary figures

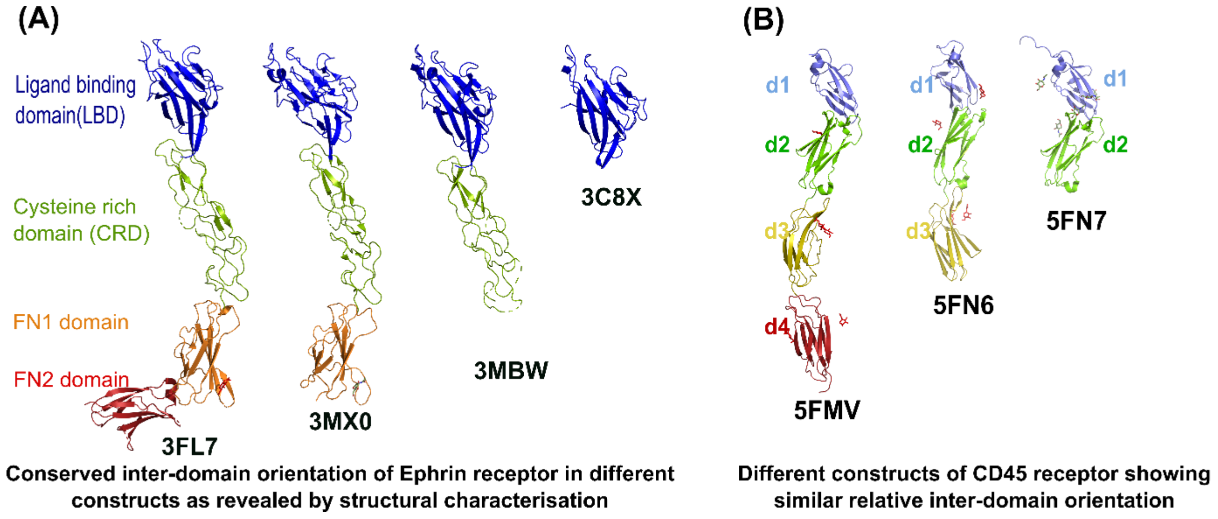


**Fig. S1** Structural comparison of different receptor constructs to show relative rigid inter-domain orientation. **(A)** Four different crystal structures of human Ephrin receptor with variable construct lengths show similar domain orientations with respect to one another (PDB Ids: 3FL7, 3MX0, 3MBW, 3C8X. **(B)** Three different crystal structures of CD45 receptor with domains d1-d4(PDB: 5FMV), d1-d3(PDB: 5FN6), d1-d2 (PDB: 5FN7) show similar rigid inter-domain relative orientation.

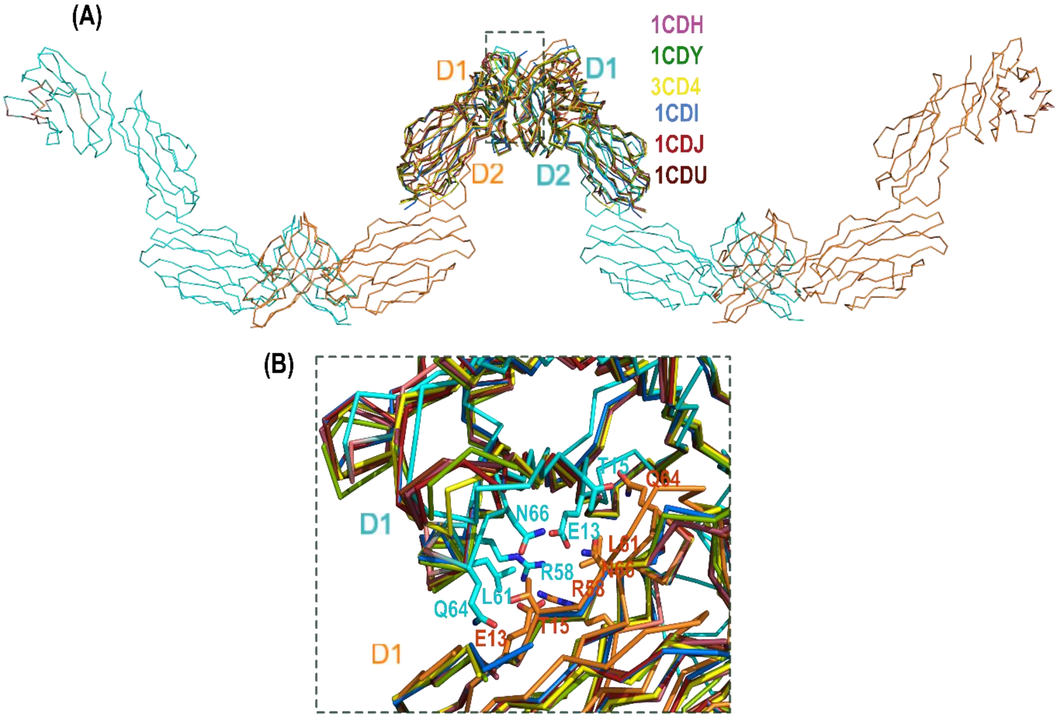


**Fig. S2** Comparison of six D1-D2 structures of CD4 with the structure of full length ectodomain of CD4 (D1-D4) **(A)** D1-D1 mediated dimers observed in six different crystal structures of CD4(D1-D2)(PDB entries: 1CDH, 1CDY, 3CD4, 1CDJ, 1CDI, 1CDU) are superposed on to the D1-D2 domains of complete ectodomain (D1-D4: 1WIO) structure. Note that the relative orientation of D1-D2 is maintained as in D1-D4 structure and D1 domains of adjacent protomers interact in a similar fashion as observed in the 1D array of D1-D4 structures **(B)** Enlarged view of D1-D1 interface showing polar interactions (shown as sticks).

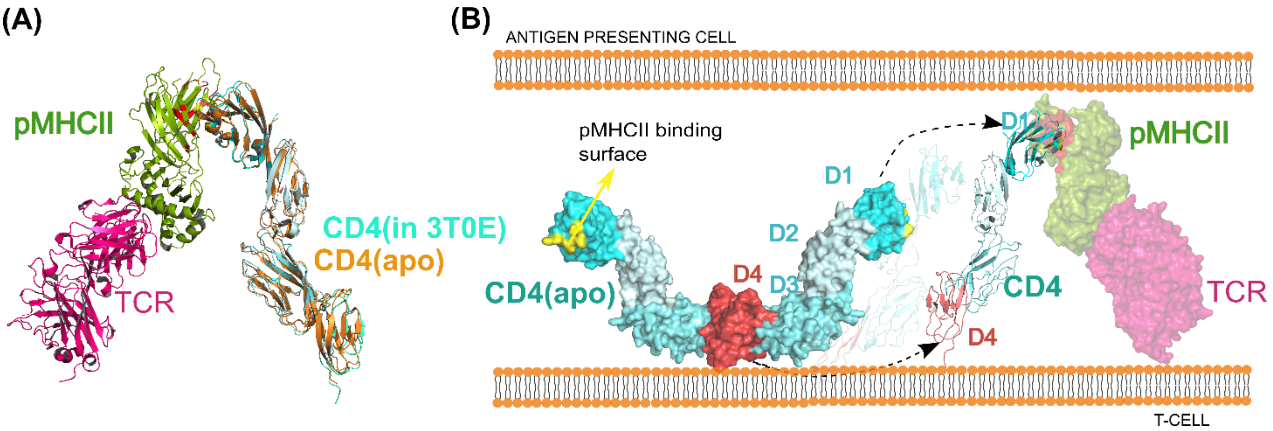


**Fig. S3** Comparison of orientation of D4 domain of CD4 with respect to membrane in apo and in complex structure: **(A)** Superposition of D1-D4 domain of apo CD4 structure (orange) on to CD4 (cyan) observed in CD4:pMHC:TCR ternary complex structure (PDB entry: 3T0E). Note that all the domains of CD4 (D1-D4) from two structures superpose well with each other, indicating overall CD4 structure in both cases are very similar **(B)**. The D4 domain (shown in brick red colour) in CD4 apo structure is seen perching on to the membrane and involved in dimerisation. Its interaction with pMHC requires reorientation of D4 domain (and probably EJ-linker) with respect to the membrane as observed in the CD4:pMHC:TCR ternary complex structure to reach its D1-mediated binding site on pMHC. The D4 domain appears to reorient itself almost perpendicular to the membrane surface, disrupting the weak dimeric interactions as well as interactions with membrane.

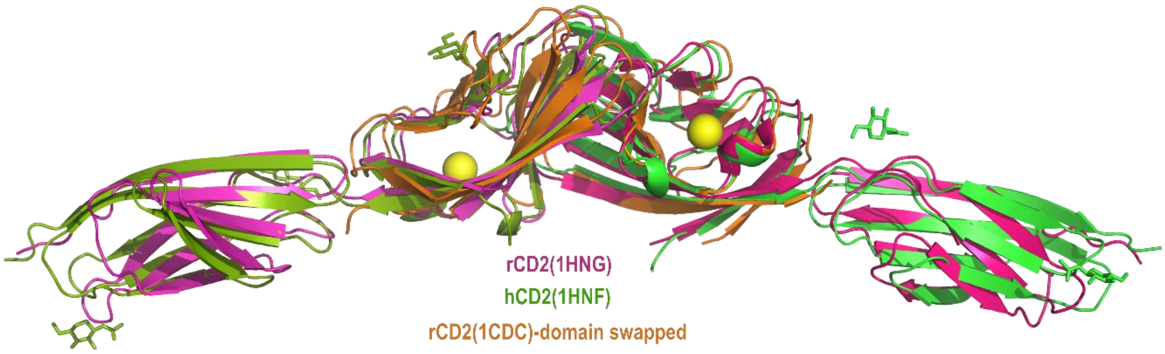


**Fig. S4** Comparison of different CD2 structures: Superposition of rCD2(PDB entry: 1HNG) (magenta), hCD2(PDB entry: 1HNF) (green) and the domain swapped structure of rCD2(PDB entry: 1CDC) (orange) reveals a similar canonical front-face to front-face interaction characteristic of the IgV:IgV interactions, although CD2 is known to be a monomer in solution.

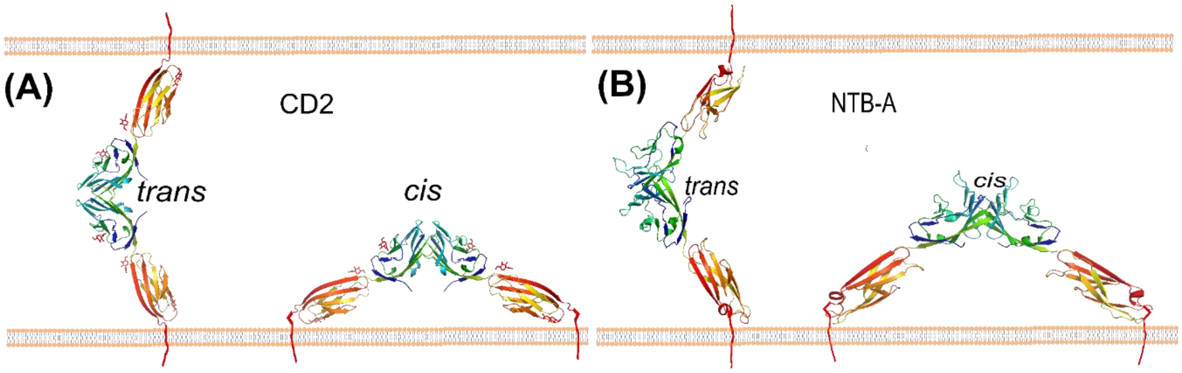


**Fig. S5** The *trans* and *cis* arrangement of CD2 and NTB-A receptors on the membrane: **(A)** hCD2 interaction from the crystal structure of PDB entry: 1HNF can be depicted as either *cis* (on same cell) or as *trans* (between molecules from opposing cells). While the *trans* interaction mimics the heterophilic interaction with CD58, homophilic *cis* interactions might be the non-signalling resting state of CD2 molecules that is observed to support 'upright' orientation. **(B)** The dimeric association as seen in the crystal structure of NTB-A (PDB entry: 2IF7) shows the protomers interacting at an angle of almost 90 degree. While such an association is often considered as *trans* association between molecules from opposing cells, from the figure it is apparent that the same can also be represented as a *cis* association of two molecules on the same cell.

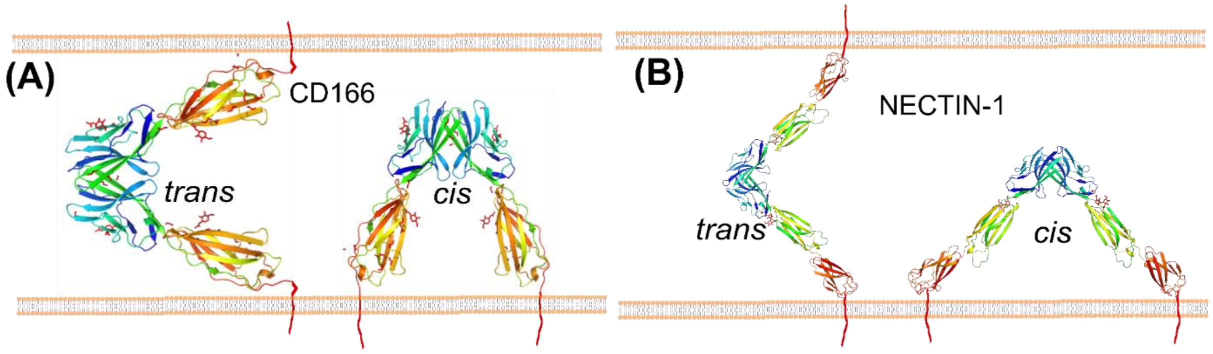


**Fig. S6** *Trans* and *cis* arrangement of CD166 and Nectin-1: **(A)** The dimeric associations as seen in the crystal structure of CD166(PDB entry: 5A2F) can be depicted as either *cis* or *trans*. **(B)** Similar to CD2, NTB-A and CD166, Nectin-1 (PDB entry: 4FMF) reveals interaction that can be represented as both *cis* and *trans*.

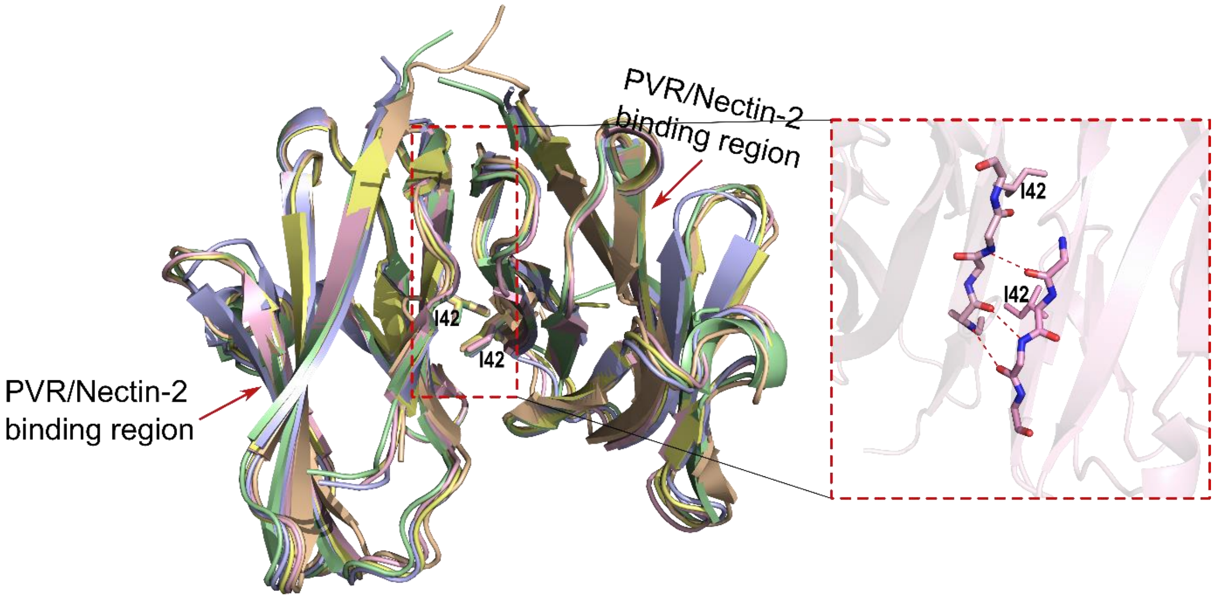


**Fig: S7** Crystal structures of TIGIT reveal *cis* oligomerisation: Five different crystal structures of TIGIT superposed on each other (PDB entries: 3Q0H, 3RQ3, 3UCR, 3UDW and 5V52. The two protomers in all the structures are connected by three H-bonds (showed in red dashes in the inset) and a hydrophobic patch with Ile42 at its centre. The mutations I42A and I42D have shown a drastic effect on receptor clustering on cell surface and signalling

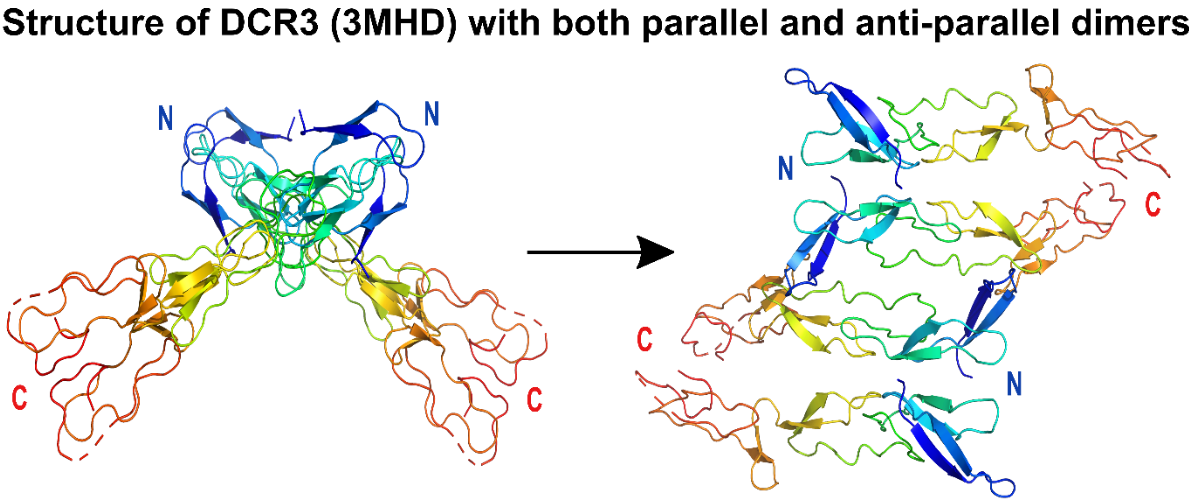


**Fig. S8** Crystal structure of DCR3(PDB entry: 3MHD) showing both parallel and anti-parallel dimers similar to TNFR1

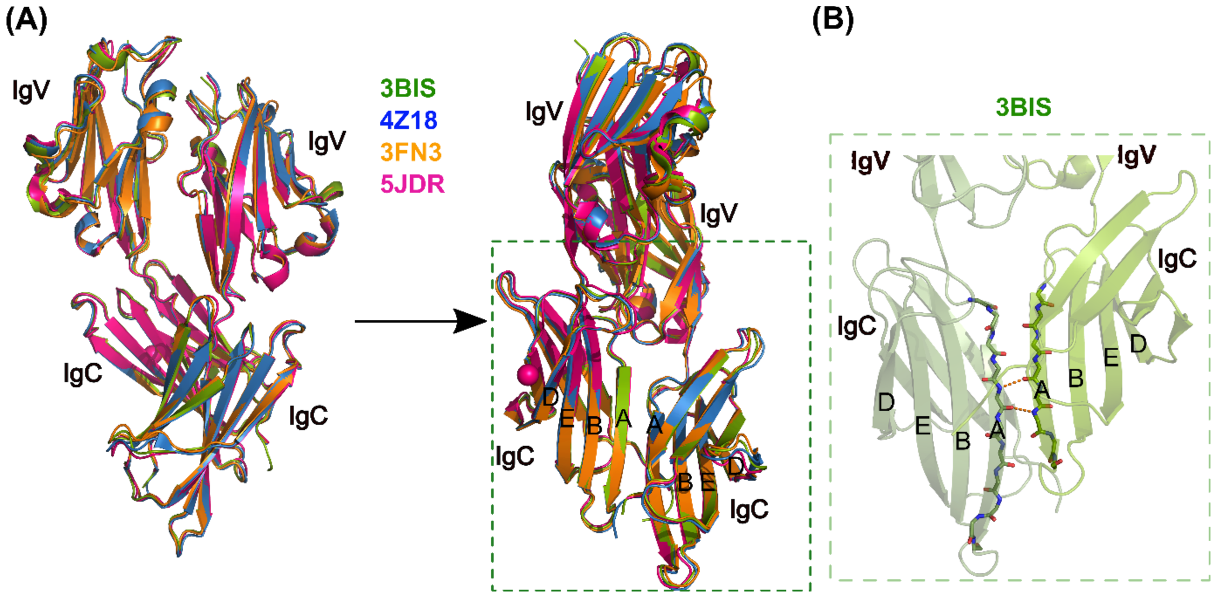


**Fig. S9** Comparison of different crystal structures of hPD-L1**(A)** Superposition of four different crystal structures of PD-L1 depicting weak dimerisation via IgC domains (PDB entries: 4Z18, 3FN3, 5JDR, 3BIS). The strands ABED of the two IgC domains fuse to form a continuous β-sheet. **(B)** Enlarged view of IgC domains of the two protomers of hPD-L1 showing fusing of β sheets. The orange dashed lines represent H-bonds between the two A-strands of the crystallographically observed weak hPD-L1 dimers.

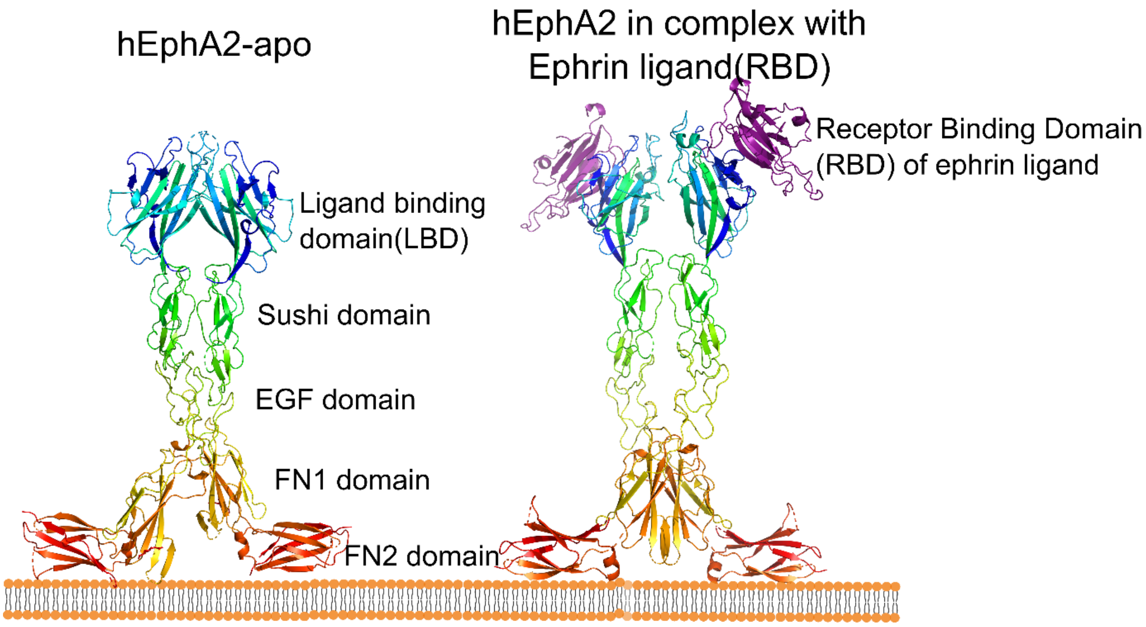


**Fig. S10** *Cis* interactions observed in the crystal structures of hEphA2 receptor: The crystal structures of hEphA2 in apo form (PDB entry: 3FL7) and in complex with receptor binding domain (RBD) of ephrin ligand (PDB entry: 2X11) shows dimeric interactions between ligand binding domains (LBD), sushi domains, EGF domains and FN1 domains of two protomers. Sushi domain and EGF domain together are also called Cysteine rich domain (CRD). FN2 domain is seen in an orientation perpendicular to the rest of the four domains of the protein. This orientation of FN2 domain resembles the D4 domain of CD4 apo structure (1WIO) as if it is perching on the membrane.
